# Temporal Mapper: transition networks in simulated and real neural dynamics

**DOI:** 10.1101/2022.07.28.501877

**Authors:** Mengsen Zhang, Samir Chowdhury, Manish Saggar

## Abstract

Characterizing large-scale dynamic organization of the brain relies on both data-driven and mechanistic modeling, which demands a low vs. high level of prior knowledge and assumptions about how constituents of the brain interact. However, the conceptual translation between the two is not straightforward. The present work aims to provide a bridge between data-driven and mechanistic modeling. We conceptualize brain dynamics as a complex landscape that is continuously modulated by internal and external changes. The modulation can induce transitions between one stable brain state (attractor) to another. Here, we provide a novel method – Temporal Mapper – built upon established tools from the field of Topological Data Analysis to retrieve the network of attractor transitions from time-series data alone. For theoretical validation, we use a biophysical network model to induce transitions in a controlled manner, which provides simulated time series equipped with a ground-truth attractor transition network. Our approach reconstructs the ground-truth transition network from simulated time-series data better than existing time-varying approaches. For empirical relevance, we apply our approach to fMRI data gathered during a continuous multitask experiment. We found that occupancy of the high-degree nodes and cycles of the transition network was significantly associated with subjects’ behavioral performance. Taken together, we provide an important first step towards integrating data-driven and mechanistic modeling of brain dynamics.

## Introduction

The brain exhibits complex dynamics (Buzsaki, 2006; Kelso, 1995). Characterizing its overall dynamic organization is a fundamental step in assessing brain functions and brain fingerprinting for healthy individuals and patients with psychiatric disorders (Saggar and Uddin, 2019). One common approach is to infer dominant “brain states” and the transitions between them from neuroimaging time series data (e.g., Cavanna et al., 2018; Li et al., 2017; Meer et al., 2020; Saggar et al., 2018; Taghia et al., 2018; Tang et al., 2012; Zalesky et al., 2014)). Such “states” and transitions can be defined by a diverse array of data-driven methods. Here we categorized a model as data-driven if it does not require additional knowledge of the brain other than the time series data recorded from it. On the other side of the spectrum, brain dynamics are often modeled by large-scale nonlinear dynamical systems models with various levels of biophysical details (Breakspear, 2017; Deco et al., 2011). Here we categorize this type of model as mechanistic, as they aim to describe the dynamical mechanism of interaction between constituents of the brain, which requires prior knowledge or assumptions about the biophysical and anatomical features of the brain in addition to the time series data measured. States and transitions discovered using data-driven methods often share conceptual appeal to nonlinear dynamics concepts such as attractors (stable states) and phase transitions. Yet, a direct link between data-driven and mechanistic modeling of the brain remains *missing*. In this work, we develop a data analysis method to represent time series data as a directed graph, whose nodes and edges could reasonably map directly to the underlying attractors and phase transitions in a nonlinear dynamic model of the brain. We first validate our method using simulated transitions and then apply the method to human fMRI data to demonstrate its empirical relevance in assessing transitions associated with cognitive task switching. This work helps build the missing link between data-driven and mechanistic modeling of complex brain dynamics. With a direct link to mechanistic models, data-driven models may better inform experimenters and clinicians of the network effect of causal perturbation (e.g., transcranial magnetic stimulation) in basic neuroscience and in the treatment of psychiatric disorders.

A signature of nonlinear brain dynamics is multistability, i.e., the coexistence of multiple stable brain activity patterns (Kelso, 2012), which may be referred to as attractors in technical terms or persistent brain states colloquially. Transitions between these brain states may occur either driven by external influences or internal dynamics. Intermittent visits to different brain states are often referred to as metastability (Tognoli and Kelso, 2014). Multistability and metastability—the existence of and the transitions between different brain states—are key elements in the mechanistic modeling of brain dynamics and functional connectivity (FC) (van den Heuvel and Hulshoff Pol, 2010). Typically, such modeling approaches use large-scale biophysical network models that also incorporate biologically informed parameters and the human structural connectome (Deco et al., 2014, 2013, 2011; Deco and Jirsa, 2012; Golos et al., 2015; Hansen et al., 2015; Zhang et al., 2022).

The mechanistic modeling of state transitions in large-scale brain dynamics was motivated by, among other things, the observations of how large-scale functional connectivity patterns vary as a function of time, i.e., the dynamic functional connectivity (dFC) (Hutchison et al., 2013; Preti et al., 2017). dFC patterns are primarily computed as correlation coefficients between time series within a sliding window. More recently, single time-frame methods (e.g., (Faskowitz et al., 2020; Zamani Esfahlani et al., 2020)) have been developed to tackle FC analysis at the finest temporal resolution and reduce the amount of data needed for stably estimating dFC patterns (Laumann et al., 2017; Leonardi and Van De Ville, 2015). Altogether, (d)FC analyses play a central role in the empirical understanding of brain dynamic organization. Abnormal FC and abnormal transitions between dFC patterns have been linked to a wide spectrum of psychiatric and neurological disorders (Barber et al., 2018; Díez-Cirarda et al., 2018; Du et al., 2021; Fox and Greicius, 2010; Garrity et al., 2007; Lui et al., 2011; Rabany et al., 2019; Saggar and Uddin, 2019).

What remains unclear is to what extent dFC patterns can be mapped to dynamical systems concepts such as attractors or stable states. With a data-driven approach, dFC patterns that repeat in time can be assigned to a relatively small number of “FC states” using, for example, clustering methods (Allen et al., 2014) or hidden Markov models (Quinn et al., 2018; Rabiner, 1989; Vidaurre et al., 2017). However, directly conceptualizing FC states as dynamical system states or attractors is not easy, especially when one needs to write down the differential equations governing the state evolution. Thus, mechanistic models of large-scale brain dynamics typically use mean-field neural activity (Cabral et al., 2017) (e.g., population firing rate, the fraction of open synaptic channels) or its derived BOLD signal (Friston et al., 2003), rather than vectorized dFC patterns, as state variables. FC states can be derived post-hoc from simulated neural dynamics (Golos et al., 2015; Hansen et al., 2015), but a *direct correspondence* between such post-hoc FC states and dynamical system attractors is yet to be demonstrated. Our recent modeling work suggests that FC patterns may be signatures of phase transitions between stable states rather than the states themselves (Zhang et al., 2022). All the above point to the need for a data-driven method to quantify stable brain states and transitions directly from time-series data and allow mapping of such states/transitions to underlying attractors and phase transitions derived from mechanistic modeling.

In the present work, we leverage existing methods of computational topology/geometry and large-scale biophysical network modeling to bridge this gap. Topological and geometrical analysis of dynamical systems traces back to the time of Poincaré (Poincaré, 1967). However, efficient computational infrastructure for generalizing such methods to higher-dimensional dynamics was not in place until recently. Morse decomposition has been used in the rigorous analysis of nonlinear dynamical systems, e.g., to represent a dynamical system as a directed graph whose nodes map to attractors (and repellers) and edges to transitions (connecting orbits) (Cummins et al., 2016; Kalies et al., 2005). However, neuroimaging data live in a very high dimensional space sparsely covered by samples, which renders many rigorous methods inapplicable. With a data-driven approach, combinatorial representations (e.g., graphs or simplicial complexes) of neural time series or FC patterns can be generated using existing topological data analysis (TDA) tools such as Mapper (Singh et al. 2007; Carlsson 2009; Saggar et al. 2018; Geniesse et al. 2022; Geniesse et al. 2019; Saggar et al. 2021) and persistent homology (Edelsbrunner and Morozov 2013; Carlsson 2009; Chazal and Michel 2021; Petri et al. 2014; Giusti et al. 2015; Billings et al. 2021). In between, there are emerging efforts to develop dynamical system-oriented TDA methods (Garland et al., 2016; Kim and Mémoli, 2021; Munch, 2013; Myers et al., 2019; Perea, 2019; Tymochko et al., 2020), some specifically supported by mechanistic models of biological dynamics (Gameiro et al., 2004; Topaz et al., 2015; Ulmer et al., 2019; Zhang et al., 2020). The present work falls under this in-between category, building on our previous work on the TDA (Geniesse et al., 2022, 2019; Saggar et al., 2018) and biophysical network modeling of large-scale brain dynamics (Zhang et al., 2022).

In the current work, our contribution is threefold. First, we introduce a novel method to extract features associated with dynamical systems (i.e., attractors and their *directed* transitions) from the observed time-series data alone. Second, to validate our approach, we develop a method to simulate a complex sequence of phase transitions in a large-scale neural dynamic model in a controlled manner. This simulated data not only provides a ground truth of the co-existing attractors in the model and their respective transitions but also allows examination of intricate but relevant nonlinear dynamic concepts such as hysteresis. Third, we apply our method to a real-world human fMRI dataset to examine the efficacy of our method in capturing states and their transitions associated with cognitive task switching from time-series data alone. Taken together, we provide a critical methodological step towards bridging the mechanistic and data-driven modeling of large-scale brain dynamics.

## Results

In this section, for larger accessibility of our results, we first provide a brief introduction to the key nonlinear dynamics concepts and intuitions. We then introduce the framework to simulate a complex sequence of phase transitions using a large-scale neural dynamic model. Finally, we present an introduction to our Temporal Mapper approach and its application to simulated as well as real-world fMRI datasets.

### 2.1 Nonlinear dynamics and the brain

Brain activities can be thought of as dynamics unfolding on a nonlinear landscape (Figure 1a). Each state in this landscape represents a pattern of activation over the whole brain. Some states are stable, which are termed attractors (Figure 1a1-3) since they attract nearby states to evolve towards them. The coexistence of multiple attractors— multistability—is a signature of the brain’s dynamic complexity (Kelso, 2012). The landscape can be shaped by a variety of intrinsic and extrinsic factors, such as external input, synaptic conductance, and structural connectivity (Zhang et al., 2022). Theoretically, these factors are often considered control parameters (Figure 1b).

**Figure 1.**
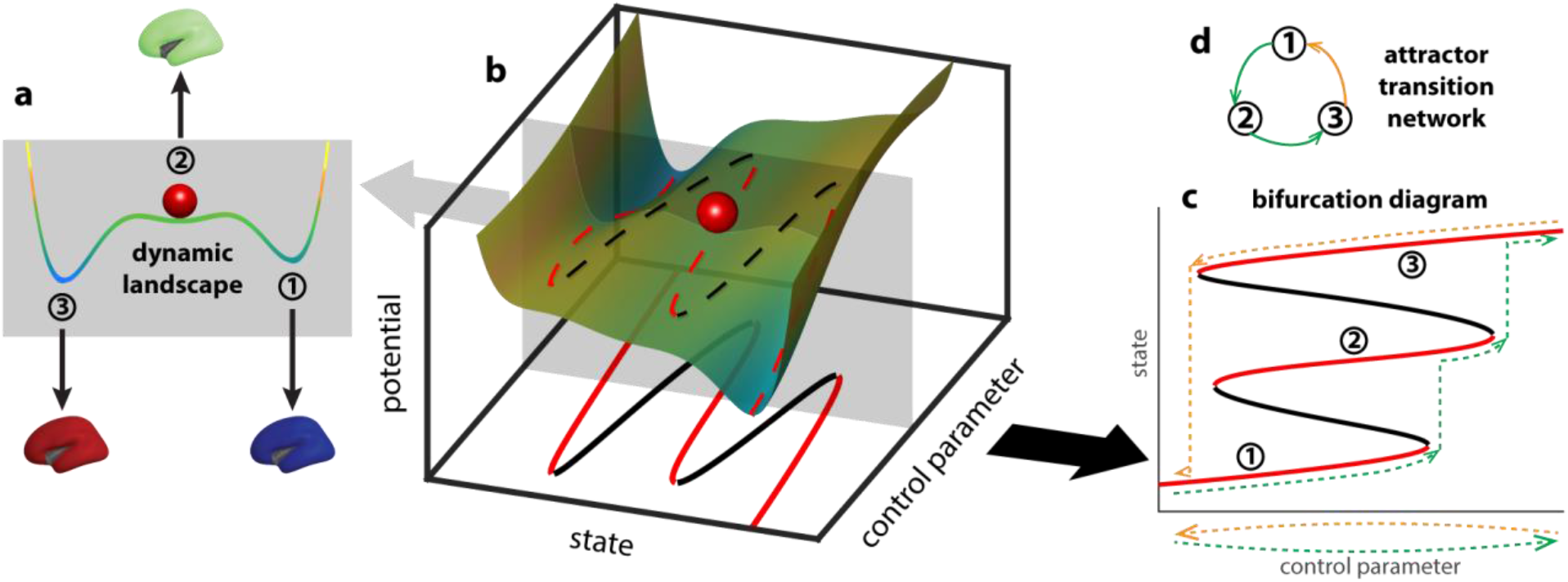
Deformation of the brain dynamic landscape induces transitions between stable brain states. A toy example of a dynamic landscape is shown as a colored curve in (a). The horizontal axis represents all possible brain states, i.e., the state space, whereas the position of the red ball represents the current brain state. States at the local minima of the landscape (a1-3) are attractors—slight perturbation of the current state (e.g., red ball) leads to relaxation back to the same state. States at the local maxima of the landscape are repellers (to the left and right of state 2, unlabeled)—slight perturbation of the state pushes the system into the basin of one of the attractors. The landscape may be deformed by continuous changes in the brain structure, physiology, or the external environment, here represented abstractly as a control parameter (b). As the landscape deforms (sliding the gray plane in b), the attractors and repellers shift continuously with it, for the most part, marked by dashed lines in red and black respectively. At critical points where an attractor and a repeller collide, there is a sudden change in the repertoire of attractors, potentially leading to a transition between attractors. The change of the landscape is commonly visualized as a bifurcation diagram (c), which keeps track of the change of attractors (red lines, 1-3) and repellers (black lines). Here “attractor” is used in a general sense, referring to both the points in the state space (the intersections between red lines and the gray plane in the bottom plane in b) and the connected components resulting from the continuous deformation of these points in the product between the state space and the parameter space (red lines in c). Due to multistability and hysteresis, the system may take different paths in the bifurcation diagram as the control parameter moves back and forth along the same line (dashed lines in c; green indicates forward paths, yellow indicates backward paths). In an even simpler form, this path dependency can be represented as a directed graph (d), denoting the sequence in which attractors are visited (color indicates forward and backward paths in c).

As the control parameter changes, the landscape deforms with it. For illustration, sliding the gray plane in Figure 1b up and down the control parameter axis changes the landscape within the plane. With sufficient multistability, changing a control parameter back and forth along the same path (Figure 1c, dashed lines below the horizontal axis) can lead to distinct paths of transitions between attractors (Figure 1c, dashed lines in the bifurcation diagram above the horizontal axis)—a feature known as *hysteresis*. Due to the asymmetry in the path of transitions, directed graphs are better suited to minimally represent the transition network (Figure 1d), where the nodes map to the attractors visited (nodes 1-3) and edges map to the transitions between attractors.

The topological complexity of this attractor transition network reflects the brain’s intrinsic complexity through its interaction with the internal or external environment. In the present work, we develop a method to retrieve such transition networks from simulated neural dynamics and human fMRI data. In the following sections, we demonstrate that the networks reconstructed from simulated data are reasonable approximations of the theoretical ground truth, and those constructed from fMRI data help predict human behavioral performance.

### 2.2 Computational framework to simulate complex sequences of phase transitions and represent them as an attractor transition network

In this subsection, we introduce the computational framework used to simulate neural dynamics. Simulations convey several advantages: (1) we can parametrically control and induce transitions between attractors, (2) we can compute the ground-truth transition network given the exact equations, and (3) we can directly compare the reconstructed network (from simulated time series alone without knowing the equations or any parameters) to the ground truth to assess the quality of reconstruction.

For simulations and computing the ground-truth transition network, we choose a biophysically informed model of human brain dynamics (Figure 2a; see Section 4.1 for details). The model describes the dynamics of a network of 66 brain regions, shown to capture functional connectivity patterns in human resting fMRI (Zhang et al., 2022) (the cartoon in Figure 2a includes 6 regions only for illustrative purposes). The model is an adaptation of the reduced Wong-Wang model (Wong and Wang, 2006; Deco et al, 2014) in the form of the Wilson-Cowan model (1972, 1973) with improved multistability (Zhang et al., 2022). Each region *i* consists of an excitatory population (E) and an inhibitory population (I), with associated state variables (*S_E_^(i)^*, *S_I_^(i)^*. Long-range connections between regions (*C_ij_* in Figure 2a) are defined by the human connectome using data from the Human Connectome Project (Civier et al., 2019; Van Essen et al., 2013) (see Methods for details). The overall strength of global interaction is modulated by an additional global coupling parameter G. We define G as the control parameter, whose dynamics (Figure 2b) modulate the shape of the underlying dynamic landscape and induce transitions between attractors through bifurcation (see bifurcation diagram Figure 2d). The simulated neural dynamics in this time-varying landscape are shown in Figure 2c. It is important to note that here we assume the control parameter G, and consequently the shape of the underlying landscape itself, is changing much slower than the state dynamics occurring within the landscape (the ball in Figure 1a can roll quickly into the valley when the landscape has barely deformed). In other words, the present conceptual framework assumes a separation of time scale between the dynamics of the control parameter (e.g. G) and intrinsic state dynamics (e.g., defined in Eq.1-2 by the time constants *τ_E_* and *τ_I_* for the excitatory and inhibitory neuronal population respectively). Physiologically, the changes in global coupling G can be interpreted as changes in the arousal level due to, for example, task demands. Recent work of Munn and colleagues (2021) suggests that cortical dynamic landscapes are modulated by ascending subcortical arousal systems mediated by the locus coeruleus (adrenergic) and the basal nucleus of Meynert (cholinergic). In particular, the locus coeruleus-mediated system promotes global integration across the cortex and reduces the energy barrier for state transitions.

**Figure 2.**
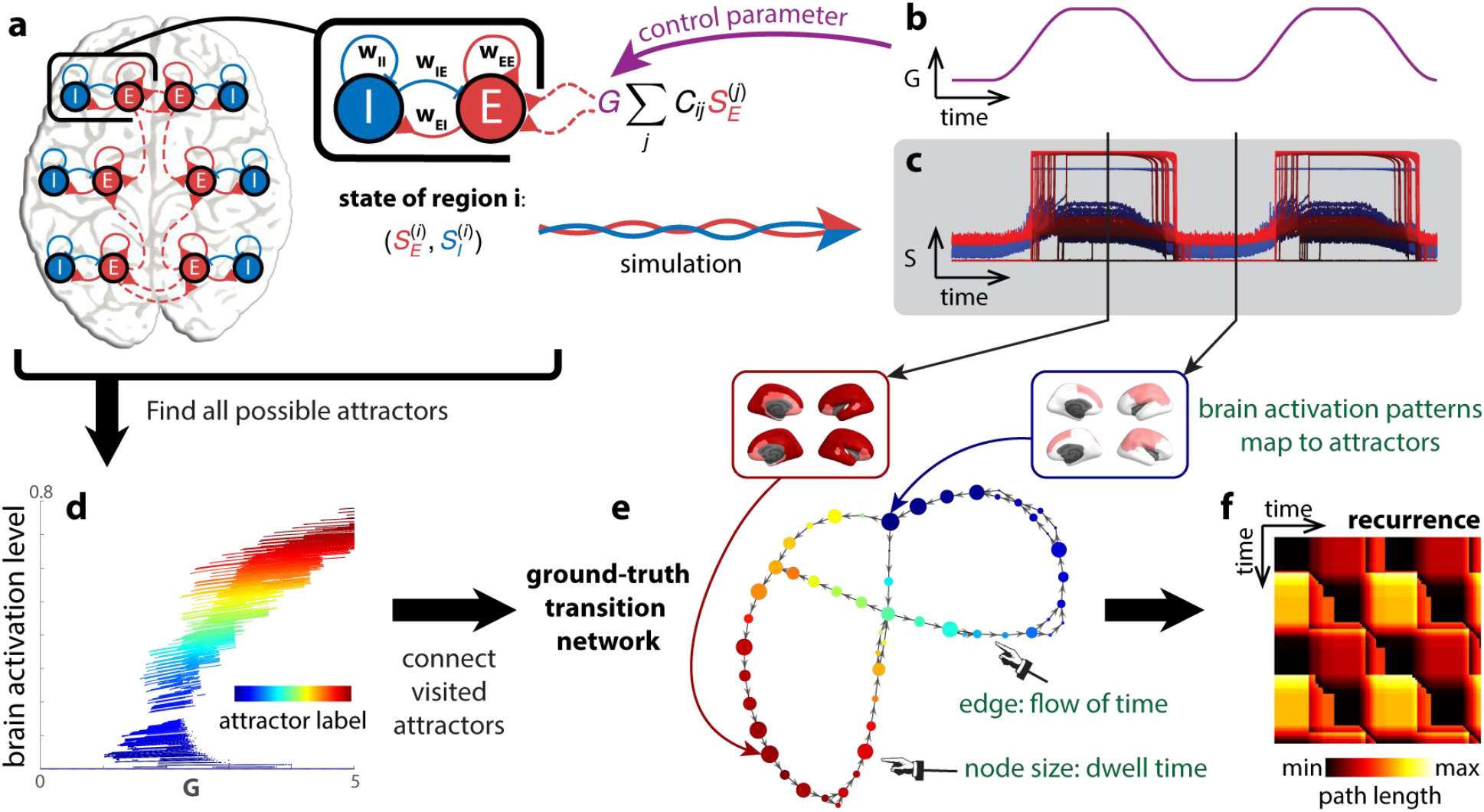
Attractor transition network for simulated neural dynamics. A biophysical network model [20] is used to describe the dynamics of the brain (a). Each brain region is modeled as a pair of excitatory (E) and inhibitory (I) populations, connected by local excitatory (*w_EE_*, *w_EI_*) and inhibitory (*w_IE_*, *w_II_*) synapses. Each region is also connected to others through long-range connections (red dashed lines). The overall strength of long-range interaction is scaled by a parameter *G*, the global coupling. To simulate neural dynamics in a changing landscape (c), *G* is varied in time (b), mimicking the rise and fall of arousal during rest and tasks. The duration of the simulation is 20 minutes. To construct a ground-truth transition network between attractors (f), fixed points of the differential equations (eq. 4-5) are computed for different levels of *G* and classified by local linear stability analysis. Fixed points classified as attractors are shown in a bifurcation diagram (d). Each attractor traces out a continuous line in a high-dimensional space—the direct product of the state space S and the parameter space *G*. These lines or attractors can be identified as clusters in S × *G*. Each time point in (b,c) is classified as the regime of one attractor in the high-dimensional space S × *G*. All visited attractors constitute the nodes of the ground-truth transition network (e), colored accordingly. A directed edge links one attractor to another if there is a transition from the former to the latter in time. To examine how dynamics unfold in time in this attractor transition network (e), we construct a recurrence plot (f) that indicates the shortest path length between any two time points (the attractors visited) in the network.

Our methodological goal is to recover the cross-attractor transitions from the simulated neural dynamics (the gating variables *S_E_^(i)^*) and the BOLD signals derived from them (down-sampled to TR=720 ms as in the Human Connectome Project (Van Essen et al., 2013)). The transitions can be encapsulated as a transition network (Figure 2e) and unfolded in time as a recurrence plot (Figure 2f). The recurrence plot depicts how far away the attractor occupied at each time point is from that of every other time point. Here, “how far away” is measured by the shortest path length from one node to another in the attractor transition network instead of the Euclidean distance between states in the original state space. The path length takes into account the underlying dynamics: two states can be far away in the state space but closely connected by transitions in the dynamical system, and conversely, two states can be close in the state space, but it could be costly to transition between each other against the underlying dynamics. The theoretical ground truth (Figure 2e,f) is constructed by assigning each time point to an attractor (Figure 2d,e) pre-computed from equations 4-5 (see methods in Section 4.2). Computation of the ground truth requires all model parameters, including the state variables *S_E_^(i)^*, *S_I_^(i)^*, and the control parameter *G* for example. As depicted, the transition network is directed to capture the “flow” of time. Further, the size of the node in the transition network represents how many sequential time points map onto that attractor, i.e., the dwell time.

We assess the efficacy of our and others’ methods by comparing the reconstructed networks (Figure 3c,d) and recurrence plots (Figure 3i,j) to the ground truth (Figure 2e,f).

**Figure 3.**
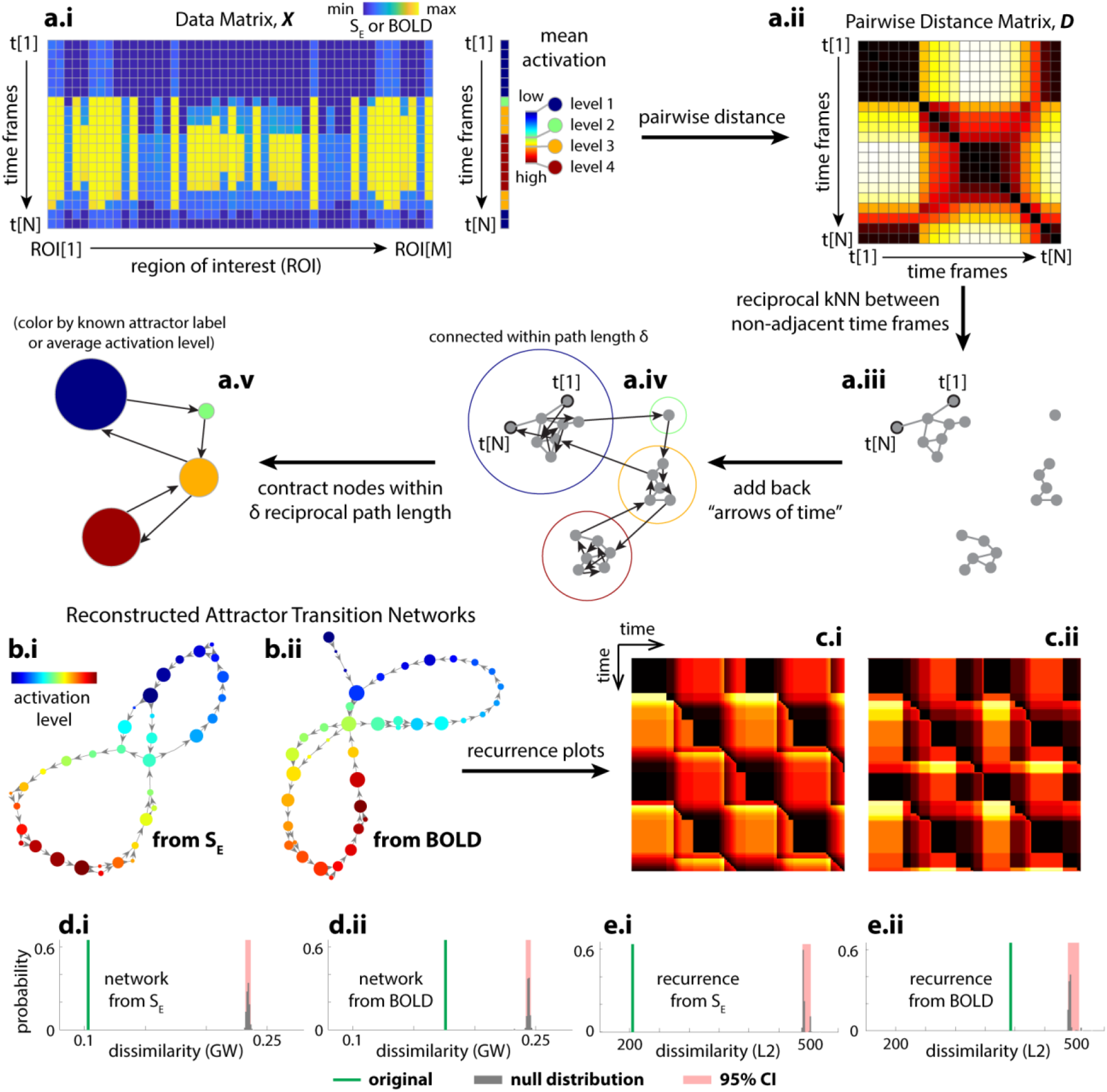
Reconstructed transition network using the Temporal Mapper approach captures theoretical ground truth. (a) shows the basic procedures of the Temporal Mapper in reconstructing attractor transition networks from time-series data. Neural time series is treated as a point cloud of N points (N time points) in an M-dimensional space (M regions of interest, ROI). As the system moves between different attractors, the activation level changes discretely. The mean activation level can be used to label each discrete state or attractor, as in Figure 2d. Pairwise distance (a.ii) between data points that are not temporally adjacent was used to construct the spatial k-nearest-neighbor (kNN) graph (a.iii). The temporal connectivity, i.e., the “arrows of time’’, is then added to the graph as directed edges (a.iv). To further compress the graph, nodes within a path length *δ* to each other are contracted to a single node in the final attractor transition network (a.v). Each node of the attractor transition network can be colored to reflect the properties of the time points associated with it (e.g., ground truth attractor labels or, when ground truth is unknown, the average brain activation level for time points associated with the node). (b.i) shows the attractor transition network reconstructed from simulated neural dynamics *S_E_* (the fraction of open synaptic channels, c.f. Figure 2c) with *k* = 16 and *δ* = 10. (b.ii) gives the attractor transition network reconstructed from the *S_E_*-derived BOLD signals with *k* = 14 and *δ* = 10, and further parameter perturbation analysis is provided in Figure S2. The node color in (b) reflects the rank of the average brain activation level for sample points associated with each node. (c.i) and (c.ii) are the recurrence plots defined for (b.i) and (b.ii) respectively. Comparing (b.i, b.ii) to Figure 2e and (c.i, c.ii) to Figure 2f, we see that the reconstructions are reasonable approximations of the ground truth. Quantitatively, we evaluate the error of approximation as the dissimilarity between the reconstructed attractor transition networks and the ground truth transition network (Gromov-Wasserstein distance, GW; green lines in d.i and d.ii) and the dissimilarity between their respective recurrence plots (L2 distance; green lines in e.i and e.ii). The reconstruction error from the original time series is significantly lower than that of randomly permuted time series (gray bars---null distribution, red area---its 95% confidence interval).

### 2.3 Temporal Mapper to reconstruct attractor transition network from time series alone

To reconstruct the attractor transition network, using only time series data, our Temporal Mapper approach first constructs a temporal version of the *k*-nearest neighbor (kNN) graph from samples in the time series (Figure 3a.i→a.iv). The time series data from multiple brain regions (Figure 3a.i) are first used to compute the pairwise distance between time points in Euclidean space (Figure 3a.ii). This distance matrix is used to determine the k-nearest neighbors for each time point. Time points that are reciprocal k-nearest neighbors (excluding temporal neighbors) are connected by edges, forming the spatial kNN graph (Figure 3a.iii). Reciprocal edges in the neighborhood graph (Figure 3a.iii) connect spatial neighbors, while directed edges connect temporal neighbors, indicating the “arrow of time” (Figure 3a.iv). Nodes that are close to each other in the neighborhood graph are contracted to a single node in its compressed version (Figure 3a.v). We consider the compressed graphs (Figure 3a.v) as reconstructed attractor transition networks. Further details of this construction are provided in Section 4.4. Visually, the reconstructions (Figure 3, b.i, c.i) are reasonable approximations of the ground truth (Figure 2e,f). This result is confirmed by quantitative comparisons against permuted time series (Figure 3d,e) and phase-randomized time series (Figure S1). The reconstruction remains robust at lower sampling rates (e.g., TR = 2s; See Supplementary Figure S11 and S12 for reconstruction accuracy drop-off rate under down-sampling) and across different realizations and integration time steps (Figure S14). See Section 4.5 for details of the graph dissimilarity measures used in these comparisons.

#### 2.3.1 The transition network, BOLD signals, and dFC reveal different facets of brain dynamics

Next, we use the simulated data to compare the dynamics in the reconstructed transition networks to its associated BOLD dynamics (from which the network is reconstructed) and dFC. Given the a priori knowledge of the ground truth, we are able to examine how different representations of the simulated time series capture different aspects of the model brain dynamics.

Dynamics in different spaces of representation (e.g., transition network, BOLD, dFC) can be compared in terms of the recurrence plots—a distance matrix that describes how far away the states at any two time points are from each other. The recurrence plot of the ground truth transition network (Figure 4a, reproduced from Figure 2g) and that of the control parameter G (Figure 4b) provide an a priori reference for comparing the reconstructed network (Figure 4c), BOLD (d), and dFC (e). In the networks (Figure 4a, c), the inter-state distance used to compute the recurrence plots is the shortest path length from one node to another. For the control parameter G and BOLD (Figure 4b, d), the distance is simply the Euclidean distance between states. For dFC (Figure 4e), the distance is the Euclidean distance between vectorized dFC matrices (Fisher-z transformed elements in the lower triangle) computed in a 30-TR sliding window. fMRI data in 20-30 TR windows have been shown to generate stable correlations (Allen et al., 2014; Hutchison et al., 2013).

**Figure 4.**
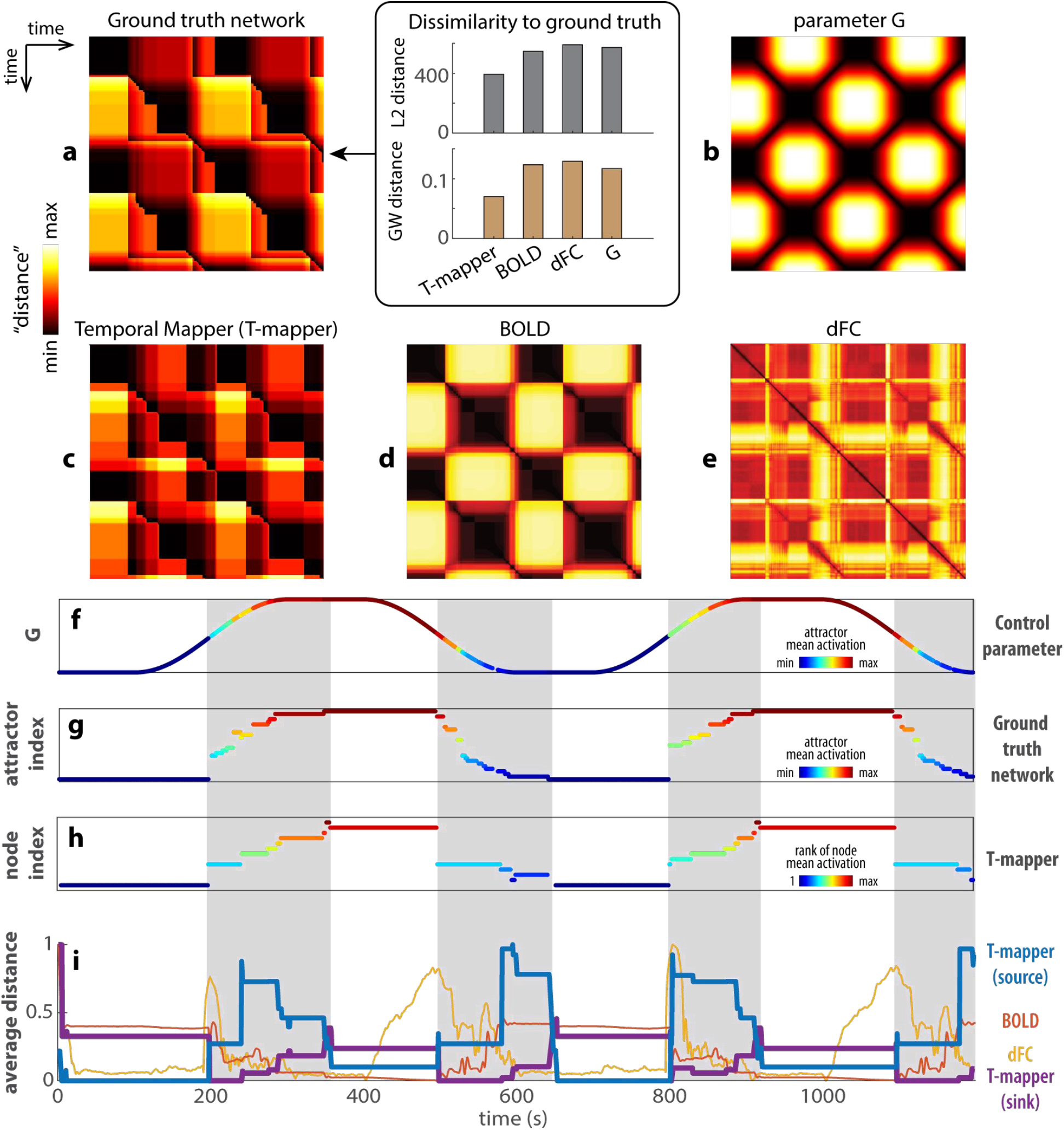
Comparisons between reconstructed transition networks, BOLD, and dFC. (a) and (b) show the recurrence plots of the ground truth transition network (a-left, reproduced from Figure 2f) and the control parameter *G* (b), respectively. They provide a basis for comparing the reconstructed transition network using the Temporal Mapper (T-mapper) (c), the corresponding BOLD signal (d), and dFC (e). The difference between the ground truth network (a) and the parameter *G* (b) reflects the organization of the underlying dynamic landscape. The greatest distinction is that the recurrent plot (a) is highly asymmetric compared to (b). The lack of symmetry in (a) reflects the path dependency and hysteresis of the underlying dynamical system. From visual inspection, the reconstructed transition network (c) is the best approximation of the ground truth network (a), especially for the asymmetric features. In contrast, the raw BOLD (d) clearly follows *G* (b) though some transitions are also visible. dFC (computed from BOLD in 30-TR windows) is neither an obvious representation of the ground truth network nor that of the parameter *G*. Quantitatively, we computed the L2- and GW distance between each recurrent plot (c, d, e, b) to the ground truth (box in a). For both measures, Temporal Mapper produces the most similar recurrence plot to ground truth, while dFC produces the most dissimilar recurrence plot. (f)-(h) compare the reconstructed network (h) more directly to the ground truth network (g) and the parameter *G* (f) in terms of the attractors visited at each point in time (only attractors that persisted greater than 5 TRs are shown). Colors in (f) and (g) reflect the attractor indices of the ground truth (y-axis of g) ordered by the global average brain activity (i.e., mean *SE*) associated with each attractor, as shown in Figure 2d. Similarly, state dynamics in the T-mapper reconstructed network (h) are ordered and colored by the global average of the simulated BOLD (rank) associated with each node. Gray areas highlight the sequence of state transitions that distinguishes nonlinear brain dynamics (g, h) from the continuous change of the control parameter (f). (i) compares the T-mapper reconstructed transition network, BOLD, and dFC by the row/column averages of the corresponding recurrence plots (c-e). Since BOLD and dFC recurrence plots are symmetrical, their row and column averages are identical (red trace for BOLD, yellow trace for dFC in i). For T-mapper reconstructed transition network, the row average is the source distance (average distance from the current state to all other states; blue trace), and the column average is the sink distance (average distance from all other states to the current state; purple trace). See text for details.

Dynamics in the ground truth and the reconstructed transition networks can also be represented as the state (attractor) dynamics shown in Figure 4g and h respectively. In this representation, we can visualize which attractors are visited during the course of time. Here, we sorted (and colored) the attractors based on their SE values (i.e., higher values mean more excitation). As evident, the dynamics extracted using the reconstructed transition network (from the Temporal Mapper approach) closely follow the dynamics of the ground truth network, indicating the nodes of the Temporal Mapper network map onto underlying dynamical attractors reasonably well. A similar visualization for BOLD and dFC can be constructed using traditional community detection approaches on the recurrence matrix (see Figure S3 for more details).

The recurrence plot for the ground truth transition network is more complex than that of the control parameter G (Figure 4a vs b)—this added complexity reflects that of the underlying dynamic landscape, which is what we aim to examine. In terms of the state dynamics, the model brain passed through a sequence of attractors and transitions in a path-dependent manner (gray areas, Figure 4g), while the control parameter undergoes continuous and reversible changes (Figure 4f, color-coded by attractor index to show the model brain can visit distinct attractors given the same level of G). Such path dependency in the ground truth transition network is indicative of multistability and hysteresis of the underlying dynamic landscape (c.f. Figure 1c). Thus, the discrete sequence of attractor transitions (gray areas, Figure 4g) is the signature of the model brain dynamics. The reconstructed transition network (Figure 4c) reasonably approximates the ground truths (Figure 4a) both in terms of the recurrence plot (Figure 4c) and the state dynamics (Figure 4h). Though some detailed transitions in the ground truth are not resolved by the reconstructed network, it is not surprising due to the low-pass filtering effect of the hemodynamic response—faster neural activity may not be recoverable from BOLD signals in principle. The recurrence plot of the simulated BOLD (Figure 4d) to a large extent follows the control parameter G, though some transitions are already visible without further state detection. Interestingly, the recurrence plot of dFC (Figure 4e) approximates neither that of the ground truth transition network nor that of the parameter G. dFC does not differentiate distinct attractors and exhibits a mixed reaction to transitions and parameter changes (see Figure S3d for how dFC states differ from attractors states in Figure S3b and BOLD states in Figure S3c).

A further comparison between the reconstructed transition network, BOLD, and dFC is shown in Figure 4i as the row averages of the corresponding recurrence plots, i.e., the average distance from the state/attractor at each time point to all other time points. This simplified representation helps us visualize the difference between dynamics in different spaces. This method was used by (Saggar et al., 2018) to examine transitions between tasks in the aforementioned continuous multitask experiment (Gonzalez-Castillo et al., 2015). To understand what information each representation captures, we separate the dynamics into two different regimes: one with a single highly persistent attractor (white areas in Figure 4g), and one with a sequence of transitions between less persistent attractors (gray area). The average distance in the reconstructed network (blue, purple curves in Figure 4i) and BOLD (red) change little in the single-attractor regime (white areas). In the same single attractor regime, the average distance of dFC (yellow curve in Figure 4i) appears to track the time derivative of the control parameter G only when G is decreasing (2nd and 4th white areas). Clearly, dFC states are not equivalent to persistent attractors. The distinction between the single-attractor and the transition regimes (white versus gray areas in Figure 4i) is best indicated by the average distance in the reconstructed network (blue, purple curve). Specifically, the persistent attractors are more source like (high sink distance, low source distance – purple above blue curve in the white areas of Figure 4i) while the sequence of transitions between less persistent attractors are more sink like (high source distance, low sink distance – purple below blue curve in the gray areas of Figure 4i). In contrast, the average distance of BOLD (Figure 4i, red) best mirrors the value of G (Figure 4f; high G is associated with low average BOLD distance and vice versa). In short, the dynamics in the reconstructed transition network most effectively separate the regimes of attractor persistence and transitions in the model brain, compared to BOLD and dFC.

### 2.4 Application of Temporal Mapper to real human fMRI dataset

Following the above theoretical analysis, we now apply the method to real human fMRI data to characterize the dynamic landscape of the human brain. We examine what features of the reconstructed transition networks are relevant to cognitive performance to demonstrate how the method developed above can be used in empirical settings. Data from 18 subjects were acquired from a continuous multi-task experiment (Gonzalez-Castillo et al., 2015). Each subject performed 8 blocks (color-coded in Figure 5g) of tasks (memory, video, math) or rest in a single 25.4-minute scan. The theoretical model used in the previous sections was designed to reflect this type of block design using the time-varying parameter G. During construction of the transition networks, we set the parameters to *k* = 5 and *δ* = 2. Justification for this choice as well as further parameter perturbation analysis is reported in Figures S9, S10.

**Figure 5.**
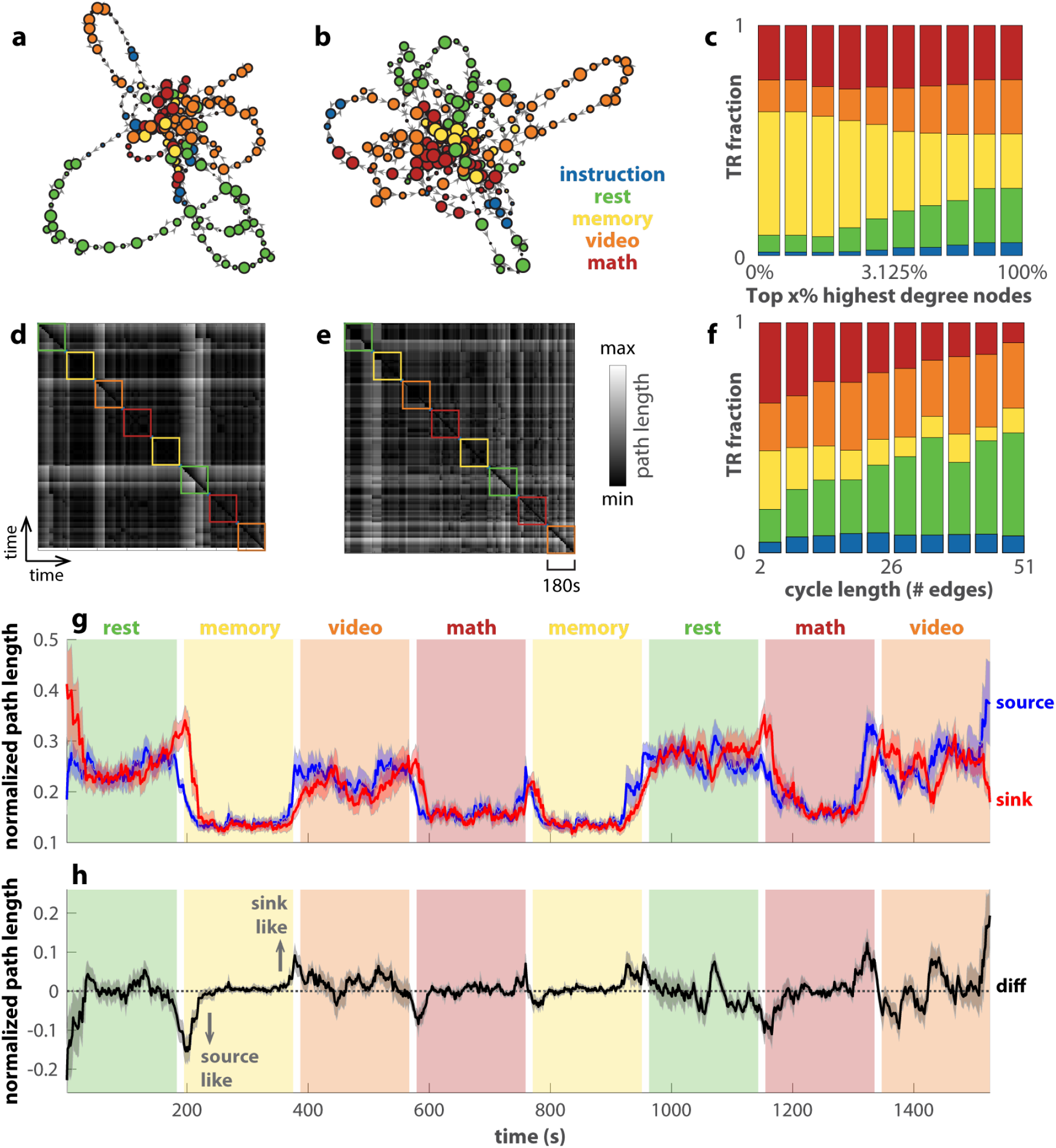
Transition networks of human fMRI data differentiate tasks and reveal transitions. (a) and (b) show the transition networks constructed from two subjects’ fMRI data in a continuous-multitask experiment as examples (Gonzalez-Castillo et al., 2015). (a) is for subject-17, among the best task performers, and (b) for subject-12, among the worst task performers. The color of each node denotes the dominant task label of the associated time points. The corresponding recurrence plots are shown in (d) and (e). (c) shows how tasks TR are distributed in the top x% highest-degree nodes of the networks across all subjects (x-axis in log scale). Memory and math clearly dominate the highest-degree nodes. In addition, (f) shows how task TRs are distributed over cycles of various lengths that pass through the top 2% of highest-degree nodes, excluding the TRs in the high-degree nodes themselves. Rest and video dominate longer cycles. (g) shows the average path length from each TR as a source to all other TRs (blue) or to each TR as a sink from all other TRs (red). The path length is normalized by the maximal distance for each subject. Solid lines show the averages across subjects; shaded areas show the corresponding standard errors. A smaller average distance indicates that the node being occupied is a better source (sink) for other nodes. The difference between the source distance and the sink distance is shown in (h). A negative (positive) number indicates that the node occupied at the time is more of a source (sink) to other nodes.

Figure 5a,d show the reconstructed transition network and the corresponding recurrence plot for a subject with good behavioral performance (top 4 in accuracy and reaction time), and Figure 5b,e for a subject with bad behavioral performance (bottom 4 in accuracy and reaction time). For both subjects, the central nodes are occupied mainly during the memory and math tasks (yellow, red nodes in Figure 5a,b) and go on excursions during the video task and rest (orange, green nodes). In aggregate across all subjects (Figure 5c), the memory task clearly dominates the highest-degree nodes (yellow bars on the left end). In contrast, rest and the video task dominate the long excursions through the highest-degree nodes (green and orange bars on the right end in Figure 5f). Later, we use this separation between the more cognitively demanding tasks (memory and math) and the relatively less demanding tasks (video and rest) over different types of nodes to predict subjects’ behavior performance (Figure 6).

**Figure 6.**
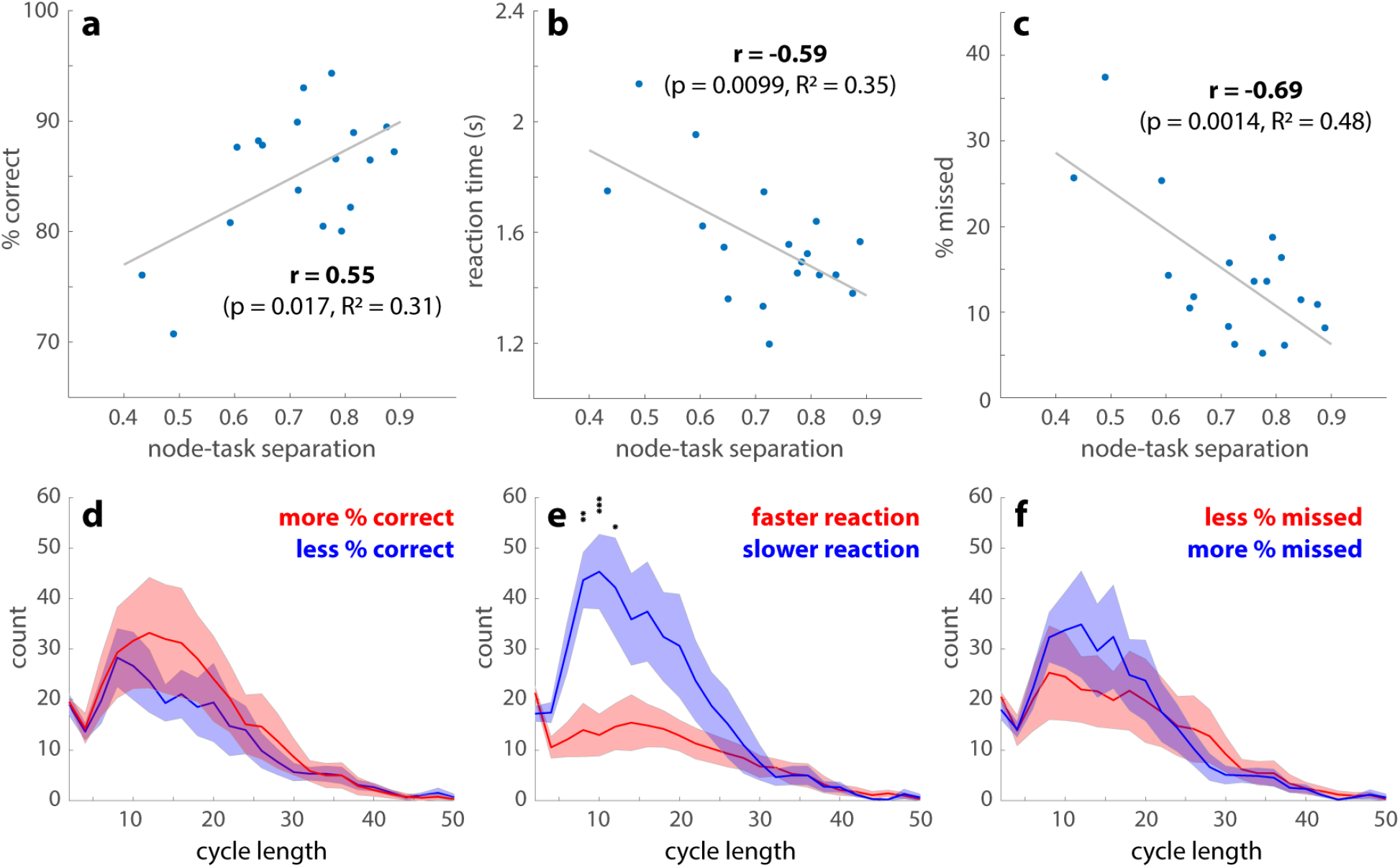
Features of transition networks predict behavioral performance. (a-c) shows how the overall task performance is associated with separations between the high-cognitive demand tasks (math and memory) and low-cognitive demand tasks (video and rest) over the transition network. The node-task separation is measured by the fraction of memory and math TRs in the top 2% highest-degree nodes of the transition networks, which also measures the preference of video and rest for low-degree nodes (c.f. Figure 5c). Subjects with a greater node-task separation have a greater percentage of correct responses (a), a faster reaction time (b), and fewer missed trials (c) across all tasks. (d-f) show the length distributions of the cycles passing through the high-degree nodes. Solid lines indicate the number of cycles at each length averaged across subjects, who are split into two groups (red vs. blue) by the median of the percentage of correct responses (d), reaction time (e), or the percentage of missed trials (f). Shaded areas indicate the corresponding standard errors. An abundance of intermediate-length cycles is associated with slower reaction time (e). There are no length-specific effects on the percentage of correct responses (d) or missed trials (f). See text for related main effects. (* p<0.05, ** p<0.01, *** p<0.001, with Tukey-HSD for multiple comparisons)

Similar to Figure 4i for the theoretical model, here we examine the dynamics in the transition networks constructed from human fMRI data in Figure 5g as the row average (source distance, blue) and column average (sink distance, red) of the recurrence plots. Both the source and sink distance clearly track the transitions between blocks. Interestingly, both math and memory are associated with a low average distance in contrast to video and rest, which suggests brain states associated with math and memory are in the more central part of the transition network. The observation is consistent with our previous findings using related methods, Mapper (Saggar et al., 2018) and NeuMapper (Geniesse et al., 2022), where the task-positive brain activation patterns were found concentrated at the core of the graph with a clear core-periphery structure. Figure 5h shows the difference between the source and sink distance. When this sink-source difference deviates from zero (black dashed line), the brain occupies a node that is either more of a source to other nodes (source-like, negative values) or more of a sink to other nodes (sink-like, positive values; see Figure S6 for example networks). The source-like or sink-like deviation is visibly more prominent near the transitions between tasks, e.g., block 2 in Figure 5h. This observation is verified statistically in Figure S7a. The source-like or sink-like deviation is also more prominent during rest and the video task than during the memory and math tasks (c.f. Figure S7b). A closer examination of the dynamics in Figure 5h reveals that the brain tends to enter the memory and math tasks through source-like nodes (downward arrow in Figure 5h) and exit the block through sink-like nodes (upward arrow in Figure 5h). This is not the case for rest and the video task. The corresponding statistical verification is shown in Figure S8. This may be due to the fact that math and memory tasks are more structured such that the brain enters the task via specific source states, while the brain can enter the resting state, for example, from any state. In short, the transition networks and recurrence plots exhibit multiple features that keep track of the task and block structure.

Figure 6 shows how features of the transition networks relate to subjects’ behavioral performance. Greater separation between cognitively demanding tasks (memory and math) and less demanding tasks (video and rest) over the highest-degree nodes predicts a higher percentage of correct responses (Figure 6a), a faster reaction time (Figure 6b) and fewer missed trials (Figure 6c). The statistical significance of the correlation coefficients is further validated against 1000 random permutations of the subject-performance correspondence (95% confidence intervals: (-0.4697, 0.5163), (-0.5123, 0.4302), (-0.5340, 0.4342) for Figure 6a,b,c respectively). The number of cycles connecting each high-degree node back to itself also exhibits behavioral relevance (Figure 6d-f). On average, a greater number of cycles is strongly associated with slower reaction time (F(1,400)=46.63, p < 10^−10^; Figure 6e). There is also a scale-specific effect—cycles around length 10 are especially predictive of slower reactions. Here the cycle length can be roughly interpreted as the number of TRs required to leave and return to a high-degree node. To a lesser extent, a greater total number of cycles is also associated with a greater percentage of correct responses (F(1,400)=4.17, p=0.014). There is no statistically significant relation between the number of cycles and the percentage of missed trials. In short, high-degree nodes and the excursions from them, i.e., cycles, are key behavioral relevant features.

## Discussion

In the present work, we propose a computational method for reconstructing attractor transition networks from neural time series, named the Temporal Mapper. The method represents a time series as a directed graph whose nodes and edges map to attractors and phase transitions in the underlying dynamical system. In particular, the method addresses the scenario where the underlying system is non-stationary or non-autonomous, as for example, when the brain is under continuously varying task demand, environmental changes, arousal levels. Using simulated brain dynamics, we demonstrate that the method provides a good approximation of the theoretical ground truth of cross-attractor transitions. Applying the method to human fMRI data, we show that the dynamics in the reconstructed networks clearly track the transitions between tasks. High-degree nodes and cycles of the network are key features that help predict human behavioral performance. Together, the theoretical and empirical analyses provide a basic theoretical and computational framework for bridging the data-driven and mechanistic modeling of brain states and transitions.

The present method builds on our previous works on characterizing neural dynamic landscape using topological data analysis (TDA). In particular, it belongs to a family of neural time series analysis tools (Geniesse et al., 2022, 2019; Saggar et al., 2021, 2018) based on Mapper (Singh et al., 2007). NeuMapper (Geniesse et al., 2022) is the closest methodological match to Temporal Mapper in that it uses a reciprocal kNN graph construction without using local low-dimensional embeddings as an intermediate step. In another variant of Mapper approach, used in (Saggar et al., 2021), an embedding step is typically utilized to examine the latent space, with any chosen dimension reduction techniques. Across the family of Mapper-based tools and related topological methods (Carlsson, 2009; Munch, 2013), time series are commonly treated as distributions of sample points in the state space. Constructing a topological representation, e.g., a graph, from such a distribution often concerns only the spatial distance between points in the state space or that of their lower-dimensional projections. The fact that these sample points are part of a time series—that there is an explicit sequential relation between sample points—is unutilized. Specifically, within the realm of fMRI data, which has been traditionally studied using time series techniques, previous applications of Mapper (Geniesse et al., 2022; Saggar et al., 2021, 2018) have focused on the geometric distribution of sample points and discarded the temporal information in the sequence of sample points. The present method is designed precisely to take advantage of this sequential information, i.e., the arrow of time. Incorporating the arrow of time better reflects the fact that the system that generates the time series, e.g., the brain, is a dynamical system. That is, the subsequent states of the system depend on the current state, and the exact nature of this dependency is the nature of the dynamical system. Restoring this temporal dependency in the construction has several consequences that we describe next.

First, the arrow of time restores the connectivity between sample points that could be far apart in the state space but tightly linked dynamically. The implication is most significant during a phase transition (vertical dashed lines in Figure 1c). At a transition, the dynamical system jumps suddenly from one attractor to another at a time scale much faster than the state dynamics within the same attractor. The combination of high velocity and even sampling in time makes consecutive sample points far apart in the state space, despite the direct dynamic dependency between them. Without the arrow of time, the increase of velocity during transitions is unaccounted for. The spatial information alone cannot determine the dynamic linkage between states of the system during the transitions, which happen to be key moments of interest.

Second, path lengths in the transition networks carry dynamical rather than purely geometrical information. The arrow of time introduces directedness into the networks. Thus, the shortest path lengths between two nodes are no longer symmetric and thus cannot be interpreted as geometric distance. The arrow of time attaches information about the underlying dynamic landscape to the state space. At an intuitive level, the shortest path from node x to node y can be considered the path of least time, or least action, within the landscape. In other words, paths in the transition networks could putatively encode actions in the state space.

Lastly, incorporating the arrow of time distinguishes the present method from common point cloud-based TDA techniques. Point-cloud data—sets of disconnected points—are fundamental substrates of topological or geometrical analysis and learning. Such analysis includes nonlinear dimension reduction techniques such as Isomap (Tenenbaum et al., 2000) or Laplacian Eigenmaps (Belkin and Niyogi, 2003), topological data analysis (TDA) methodologies such as persistent homology (Carlsson, 2009; Edelsbrunner and Morozov, 2013), and deep learning (Qi et al., 2017). With the points first connected to their temporal neighbors, the present method operates on, in essence, a discretized version of a curve with a direction defined on it—naturally depicting a trajectory of an autonomous, deterministic dynamical system. Constructing a spatiotemporal neighborhood graph (Figure 3b) is thus folding a directed curve rather than “connecting dots”. An exposition of the mathematical consequences of the construction is beyond the scope of the present work but worthy of further study, especially with regard to its behavior as the sampling rate of the time series approaches infinity and when multiple trajectories are included in the construction.

One may find the transition networks in the present work reminiscent of hidden Markov models (HMM) for detecting discrete states in brain dynamics and transition probabilities between states (Baker et al., 2014; Meer et al., 2020; Ou et al., 2013; Rezek and Roberts, 2005; Vidaurre et al., 2017). A key distinction is that the number of discrete states in an HMM is set a priori, while its analog in the present method—the number of attractors visited—is data-driven. The dynamic landscape of the human brain can be highly complex and sensitive to the organization of the large-scale structural connectome (Zhang et al., 2022). There is no a priori way to determine how many attractors may be visited during tasks and rest. Moreover, the dynamic landscape of each subject is shaped by the structural connectivity in each subject’s own brain. It cannot be assumed that the same number of attractors would be visited across subjects. Thus, the present method presents a flexible framework that requires fewer assumptions about the underlying dynamical system. For example, the Markov property for HMM (dependency on present state only) may not be satisfied in non-autonomous dynamical systems as conceptualized in the present work (see Figure S13 for an application of HMM on the simulated data), while the Temporal Mapper does not require the Markov property. Apart from statistical inference methods such as the HMM, topological representations of attractor networks also figure in differential equation-oriented state-space decomposition methods, e.g., based on the Conley Index Theory (Ban and Kalies, 2006; Kalies et al., 2005). In comparison to such mathematically rigorous approaches, the present method better accommodates empirical applications where the state space is often sparsely covered by data and the underlying dynamical system is changing with time. Conceptually, the attractor transition networks in the present study should be thought of not as representing the structure of an autonomous dynamical system, but rather the complex interaction between the environment and the brain dynamic landscape.

Our flexible, individualized construction of transition networks comes with a cost—there lacks a direct correspondence between different networks. An advantage of an HMM constructed from data concatenated across subjects (c.f. Meer et al., 2020) is that the model comes with a fixed number of states across all subjects, with direct one-to-one correspondence. In contrast, the correspondence problem for the present method is data-driven and needs to be solved separately. For example, attractor transition networks for different subjects (Figure 5a,b) contain different numbers of nodes (attractors) and edges (transitions). How nodes and edges in Figure 5a map to those in Figure 5b is not obvious. Even when comparing attractor transition networks with the same number of nodes, obtaining a correspondence is equivalent to solving an instance of the Graph Matching problem, which is a hard, fundamental challenge in graph processing (Umeyama, 1988; Zaslavskiy et al., 2009). In the present work, the correspondence between networks is made using techniques from optimal transport theory, specifically the use of Gromov-Wasserstein (GW) matchings between networks (Figure 3e,g) which can provide approximate graph matching solutions even for graphs with different numbers of nodes. GW matching does not require a temporal correspondence or any a priori knowledge of how the networks are constructed, and can thus be used in a broad context. For example, for resting-state fMRI, there is no a priori way to map the transition network constructed from one recording session to that of another. GW matchings provide a solution to compare a subject’s resting transition networks in different sessions to examine the stability of the brain dynamic landscape, or to compare transition networks of healthy subjects to that of the psychiatric patients to examine the dynamic basis of the disorder. We therefore introduce this tool to the neuroimaging community with broader future applications in mind.

The present work also brings attention to the interpretation of dFC states. It is shown in Figure 4e,i and Figure S3d that dFC states are not equivalent to attractors in the brain dynamics: dFC is in part sensitive to the transitions (Figure 4i gray areas) and in part to the change in the control parameter without inducing any phase transition (Figure 4e white areas). It has been proposed that resting-state FC reflects cross-attractor transitions (Zhang et al., 2022). While dFC-based clustering can differentiate tasks very well (Gonzalez-Castillo et al., 2019), this differentiation may not be interpreted as the brain occupying distinct attractors during different tasks, but rather, these tasks involve different transitions and environment-driven correlations. Complementing existing dFC-based approaches, the attractor transition networks incorporate information of the underlying dynamical system and provide a channel to connect data-driven representations to dynamical systems concepts. Future work may further explore the relation between attractor transition networks and dFC by using single-frame techniques for FC quantification (Esfahlani et al., 2020; Faskowitz et al., 2020).

In application to the human fMRI data, we provide a series of analyses comparable to those of an earlier work (Saggar et al., 2018), where undirected networks were constructed using Mapper (Singh et al., 2007) from the same dataset (Gonzalez-Castillo et al., 2015). Saggar et al (Saggar et al., 2018) observed that the core of the networks was dominated by cognitively demanding tasks, such as memory, math, and to a lesser extent, video tasks, while the peripheries were dominated by rest. In the same vein, we observe that the highest-degree nodes are dominated by memory and math (Figure 5c), and that the level of dominance predicts behavioral performance (Figure 6a-c). Note that due to the compression step in the present construction (Figure 3b→c), tightly connected nodes in the neighborhood graph (Figure 3b) are contracted to a single node in the final attractor transition network (Figure 3c). It is reasonable to assume that tightly connected nodes that serve as the core for periphery nodes in the neighborhood graph would become a high-degree node in the compressed graph. Thus, the high-degree nodes in the present work may be thought of as a loose counterpart to the core of the Mapper-based construction (Saggar et al., 2018).

The periphery structures of interest in the present construction are the cycles that connect the high-degree nodes back to themselves. For the human fMRI data, long cycles are dominated by rest and the video task (Figure 5f), analogous to the Mapper-based result (Saggar et al., 2018) that periphery nodes are dominated by rest. Importantly, the present cycle-based analysis of the periphery structure allows us to examine recurrent dynamics of different duration (reflected as cycle length, Figure 5f) and identify the behavioral relevant time scales (Figure 6d-f). Interestingly, slower reaction time during tasks is associated with an excess of intermediate but not long cycles (Figure 6e). This suggests that intermediate cycles putatively represent the stray path that the brain took, resulting in slower reaction times. Interestingly, a greater number of cycles predicts higher accuracy, which, combined with the reaction time results, may reflect a speed-accuracy trade-off. It provides an example of process-based analysis of brain dynamics afforded by the present, dynamics-minded, construction. From a more technical viewpoint, the spectrum of cycle length (Figure 6d-f) is tightly connected to the multiscale nature of the attractor transition network. The Temporal Mapper naturally admits multiscale construction via the compression distance δ (Figure 3a.iv). That is, one can consider the attractor transition network as a sequence of networks at different scales δ (see Figure S9 for example) instead of a single graph. At a specific scale, any smaller recurrent processes, i.e., smaller cycles formed by a few recurring attractors, will be contracted into a single node. Thus, the spectrum of cycle length indicates at which scales the graph can be further compressed. Further method development and empirical testing are required to better take advantage of the multiscale information in the Temporal Mapper. In addition, unfolding the dynamics in time as the average distance in the transition network tracks the transition between tasks (Figure 5g), comparable to Mapper-based results (Saggar et al., 2018). In the present work, the directed nature of the attractor transition networks introduces additional information—the direction of least action between states. Some nodes have a characteristic direction as a sink or a source (positive or negative deviation in Figure 5h), serving as entry and exit for the cognitively demanding tasks (Figure S8). Note that the sink-/source-ness of the associated brain activity patterns may not be absolute, as it is possible for the sink-/source-ness to depend on the specific design of the experiment (e.g., what types of tasks are included in the session and the time allocation for each task). Further studies are necessary to elucidate the interpretation of sink/sourceness in a more general context, which will require applying the Temporal Mapper to a wider range of datasets. Nevertheless, the present study demonstrates that the directedness introduced by the arrow time is task-specific. Future work may further explore the cognitive correlates of different directed features of the graph, including the sink/sourceness of the nodes. Moreover, this directedness may be useful for designing neural stimulation protocols to more effectively perturb the brain into desirable states.

In conclusion, we propose a computational method for constructing attractor transition networks from simulated and empirical time series of brain dynamics. Complementing existing geometrical and statistical approaches to characterizing brain dynamics, the present work aims to provide a channel of correspondence between data-driven topological modeling and mechanistic modeling. Incorporating time in the construction of spatiotemporal neighborhoods, paths in the attractor transition networks encode the action of the underlying dynamical systems. The method is both validated using a biophysical network model of the brain and shown to reveal behavioral and cognitive relevant features in human fMRI data. The present work serves as a starting point for dynamical theory-driven topological analysis of brain dynamics. Future work will compare the present state and transition detection methods more extensively to existing community detection methods and further validate the method using consortium-sized data (Gordon et al., 2017; Saggar et al., 2021; Smith et al., 2013).

## Materials and Methods

### 4.1 Biophysical network model of the human brain

The theoretical components of the present work are based on a biophysical network model of large-scale brain dynamics (Zhang et al., 2022), which is an variant of the reduced Wong-Wang model (Wong and Wang, 2006; Deco et al, 2013, 2014). The model is a Wilson-Cowan type model (Wilson and Cowan, 1972, 1973). The whole brain is modeled in terms of the mean-field activity of neuronal populations in each brain region (Figure 2a). Each model region contains a pair of excitatory (E) and inhibitory (I) populations, whose activity is described by the *local model* (Figure 2a, right box) in terms of the state variables *S_E_* and *S_I_*:

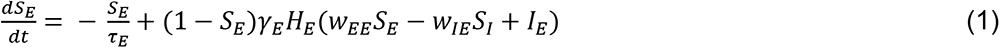

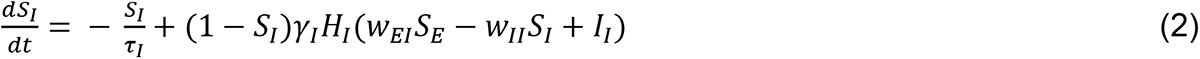

*S_E_* and *S_I_* indicate the fraction of open synaptic channels in their respective populations, referred to as the gating variables. Through local connections (*w*’s), the excitatory population excites itself with strength *w_EE_* and the inhibitory population with strength *w_EI_*, while the inhibitory population inhibits itself with strength *w*_II_ and the excitatory population with strength *w_IE_*. Each population can also receive input from outside of this region, denoted as *I_E_* and *I_I_*. The activity of each population has a natural decay time of *τ_E_* and *τ_I_* respectively. Each population’s activity tends to increase with the fraction of closed channels (1 − *S_p_*) and the population firing rate (*H_p_*), scaled by a factor *γ_p_* for *p* ∈ {*E, I*}. *H_E_* and *H_I_* are transfer functions that map synaptic current input to population firing rate of the excitatory and the inhibitory population respectively. In particular, they are sigmoidal functions of the form

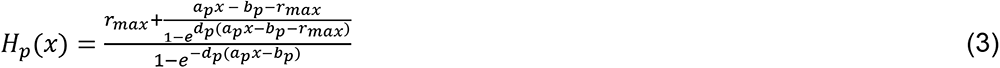

whose output increases with input monotonically and saturates at *r_max_*---the maximal firing rate limited by the absolute refractory period of neurons. The specific shape of each transfer function is determined by three additional parameters *a_p_*, *b_p_*, and *d_p_* (*a_p_* and *b_p_* determine the location and slope of the near-linear segment in the middle; *d_p_* determines the smoothness of the corners bordering the said near-linear segment). This transfer function is converted from Wong and Wang’s original formulation (Abbott and Chance, 2005; Wong and Wang, 2006) (a soft rectifier function, adopted into large-scale biophysical network models of the brain by Deco et al, 2013, 2014) into a sigmoidal form, while retaining the original value of parameters *a_p_*, *b_p_*, and *d_p_* (shown in Table 1). The parameters were chosen to approximate the average response of a population of spiking pyramidal cells (*p* = *E*) and interneurons (*p* = *I*) respectively, incorporating physiologically plausible parameters (Wang, 2002; Wong and Wang, 2006).

**Table 1.**
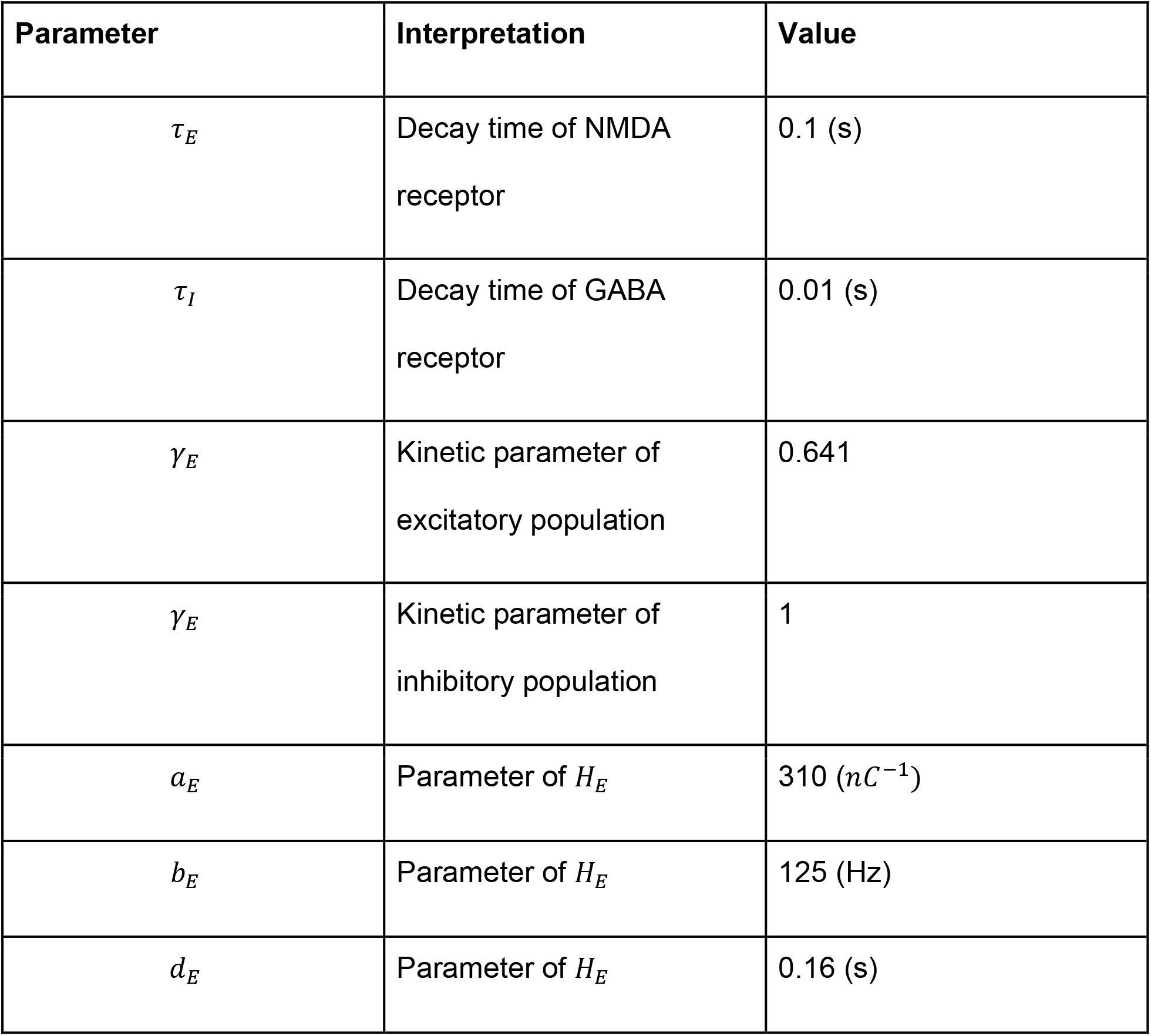
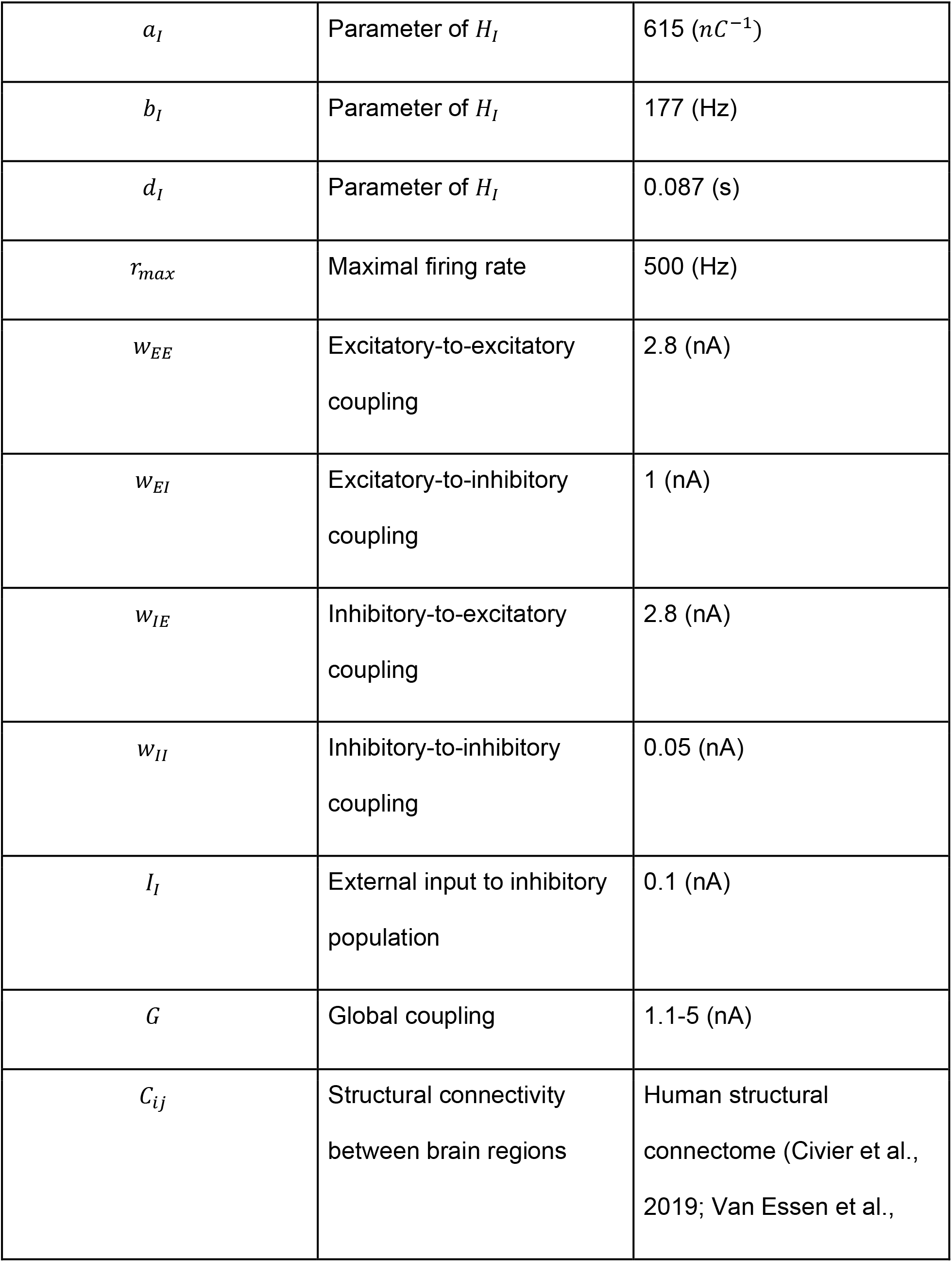
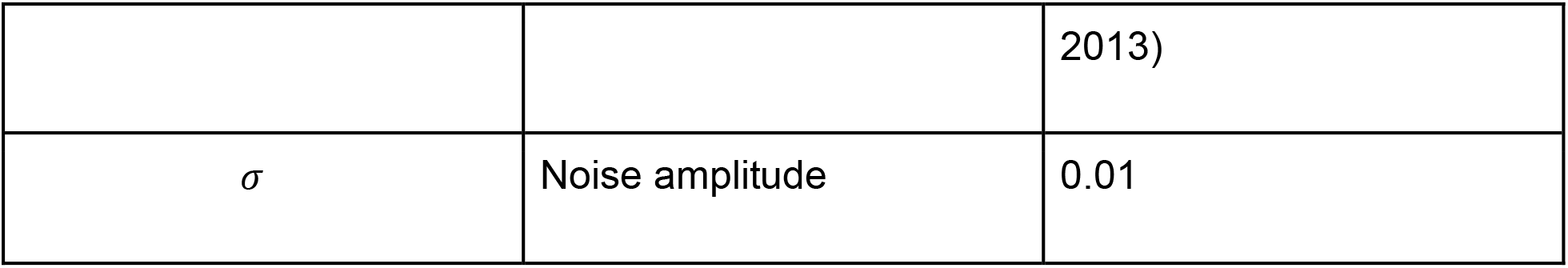
The interpretation and value of model parameters. Here we summarize the parameters used in the global model (equation 4-5). Critical parameters are inherited from (Wong and Wang, 2006) to maintain biological plausibility.

| Parameter | Interpretation | Value |
| --- | --- | --- |
| $\tau_E$ | Decay time of NMDA receptor | 0.1 (s) |
| $\tau_I$ | Decay time of GABA receptor | 0.01 (s) |
| $\gamma_E$ | Kinetic parameter of excitatory population | 0.641 |
| $\gamma_E$ | Kinetic parameter of inhibitory population | 1 |
| $a_E$ | Parameter of $H_E$ | 310 ($nC^{-1}$) |
| $b_E$ | Parameter of $H_E$ | 125 (Hz) |
| $d_E$ | Parameter of $H_E$ | 0.16 (s) |
| $a_I$ | Parameter of $H_I$ | 615 ($nC^{-1}$) |
| $b_I$ | Parameter of $H_I$ | 177 (Hz) |
| $d_I$ | Parameter of $H_I$ | 0.087 (s) |
| $r_{max}$ | Maximal firing rate | 500 (Hz) |
| $w_{EE}$ | Excitatory-to-excitatory coupling | 2.8 (nA) |
| $w_{EI}$ | Excitatory-to-inhibitory coupling | 1 (nA) |
| $w_{IE}$ | Inhibitory-to-excitatory coupling | 2.8 (nA) |
| $w_{II}$ | Inhibitory-to-inhibitory coupling | 0.05 (nA) |
| $I_I$ | External input to inhibitory population | 0.1 (nA) |
| $G$ | Global coupling | 1.1-5 (nA) |
| $C_{ij}$ | Structural connectivity between brain regions | Human structural connectome (Civier et al., 2019; Van Essen et al., |
|  |  | 2013) |
| $\sigma$ | Noise amplitude | 0.01 |

Connecting the local models into a global network (Figure 2a, left, dashed lines) gives us the *global model*:

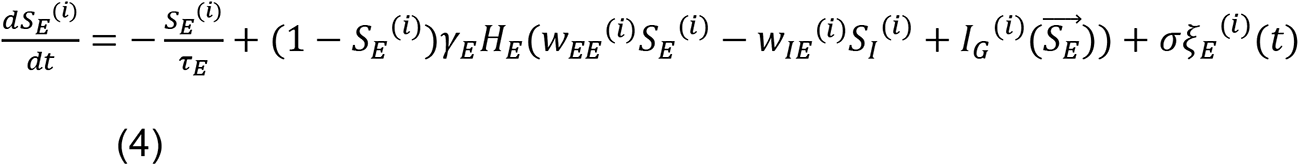

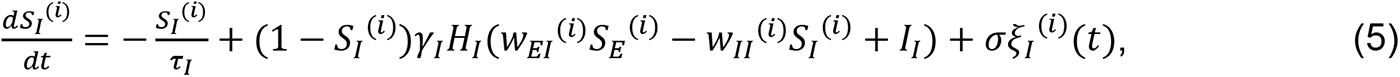

where *S_E_*^(*i*)^ and *S_I_*^(*i*)^ are the synaptic gating variable of the excitatory and the inhibitory population of the *i-*th brain region respectively, and *ξ^(*i*)^* is a noise term scaled to an amplitude *σ*. For computing the fixed points, *σ* = 0; for numeric simulations, *σ* = 0.01 following Zhang et al (2022). The state of all excitatory populations is denoted as a vector 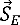 the *i-*th element of which is *S_E_*^(*i*)^. The global input to the *i-*th brain region depends on both its connectivity with, and the ongoing state of, other brain regions,

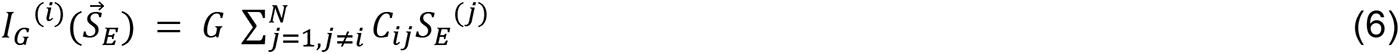

where *N* denotes the total number of brain areas, *C_ij_* ≥ 0 the long-range structural connectivity from the *j*-th to the *i*-th brain region and *G* is a global coupling parameter that controls the overall level of interaction across brain regions. Since *C_ij_* is only intended to represent long-range connectivity, we let *C_ij_* = 0 for any *i* = *j* to preclude recurrent connections. For the effects of *G*and *C_ij_* to be independently comparable, here we impose a normalization condition on the matrix norm:

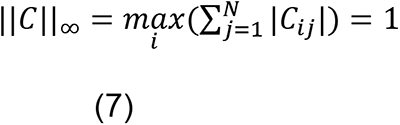

In the present work, nodes of the global network correspond to anatomical regions in the human brain based on a 66-region parcellation used in (Deco et al., 2013; Hagmann et al., 2008) (Table S1); the weight of edges reflects the strength of long-range structural connectivity between brain regions, estimated using structural data from the Human Connectome Project (Civier et al., 2019; Van Essen et al., 2013). Specific parameter values are given in Table 1.

### 4.2 Computation of the attractor repertoire and ground-truth transition network

Fixed points of the global model (equations 4-6) are computed using *fsolve* in MATLAB for each parameter *G* (in 0.01 increments). A subset of the fixed points is further classified as attractors based on local stability analysis using the Jacobian matrix and simulated perturbation. These attractors are visualized in Figure 2d as a bifurcation diagram (see (Zhang et al., 2022) for further computational details). Attractors that continuously map to each other under the variation of parameter *G* are considered qualitatively equivalent. Thus, in the present work, the term “attractor” is used in a broad sense, referring to the connected components of (narrow-sense) attractors in the product of the state space and the parameter space. Computationally, single-linkage clustering is used to obtain these connected components. The set of all connected components or clusters constitutes the attractor repertoire, providing a compact description of the model dynamic landscape (Zhang et al., 2022).

The attractor repertoire further provides a skeleton to convert any simulated times series of the state variables (within the appropriate parameter range) into the corresponding symbolic dynamics. Here symbolic dynamics refers to a sequence of symbols, where each symbol represents an attractor, and the sequence represents the order in which the attractors are visited in time. Computationally, each time point in the simulated time series is assigned to an attractor using nearest-neighbor classification in the product of the state space and the parameter space, given the pre-computed attractor repertoire. The resulting symbolic dynamics is used to construct the ground-truth attractor transition network (Figure 2f). Nodes of the network are unique symbols appearing in the sequence. There is a link from node *i* to node *j* if the corresponding symbol *i* appears immediately before symbol *j* in the sequence. Each node is equipped with a probability measure proportional to the dwell time at the corresponding attractor (i.e., node size in Figure 2f).

### 4.3 Simulated brain activities and BOLD signals

We simulate time series of brain activities and the derived BOLD signals as substrates for data-driven construction of attractor transition networks (Section 4.4). The neural activity of each model brain region *i* is represented by the local excitatory population activity *S_E_*^(*i*)^ (equations 4-6), simulated using the Heun stochastic integration scheme with a 0.001 s time step.

The derived BOLD (Blood-oxygen-level-dependent) activities are computed using the Balloon-Windkessel model (Buxton et al., 1998; Friston et al., 2003, 2000; Mandeville et al., 1999),

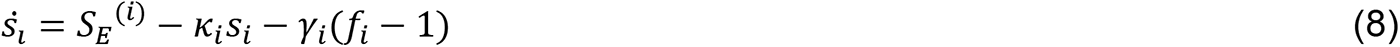

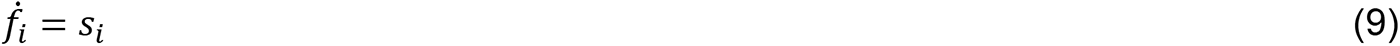

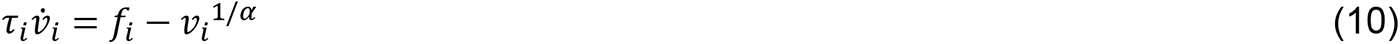

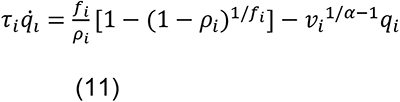

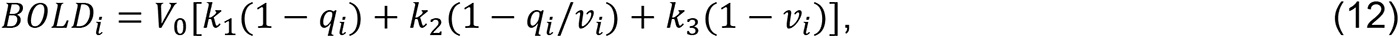

where the interpretation and values of the parameters are given in Table 2. The initial condition is:

**Table 2.** Parameters of the Balloon-Windkessel model of BOLD activity, obtained from (Friston et al., 2003).

| Parameter | Interpretation | Value |
| --- | --- | --- |
| $s_i$ | Vasodilatory signal | Variable |
| $f_i$ | Blood inflow | Variable |
| $v_i$ | Blood volume | Variable |
| $q_i$ | Deoxyhemoglobin content | Variable |
| $\kappa_i$ | Rate of signal decay | $0.65(s^{-1})$ |
| $\gamma_i$ | Rate of flow-dependent elimination | $0.41(s^{-1})$ |
| $\tau_i$ | Hemodynamic transit time | $0.98 (s)$ |
| $\alpha$ | Grubb's exponent | 0.32 |
| $\rho$ | Resting oxygen extraction fraction | 0.34 |
| $V_0$ | Resting blood volume fraction | 0.02 |
| $k_1$ | BOLD weight parameter | $7\rho_i$ |
| $k_2$ | BOLD weight parameter | 2 |
| $k_3$ | BOLD weight parameter | $2\rho_i - 0.2$ |

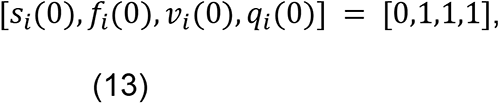

which is a hemodynamic equilibrium state without neural activity. The BOLD dynamics are passively driven by neural activity *S_E_*^(*i*)^, simulated using the Euler method with an integration time step of 0.001 *s*.

### 4.4 Temporal Mapper – construction of transition networks

The fundamental backbones of our transition networks are the k-nearest-neighbor graphs (kNNG) that often appear in dimension reduction techniques (Van Der Maaten et al., 2009). Before introducing this construction, we first set up some preliminaries.

Given a *d*-dimensional vector *v* and *p* ≥ 1, the *p*-norm of *v* is defined by writing 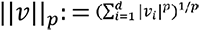. Unless specified otherwise, we will use *p* = 2 throughout this work, i.e. the Euclidean norm. Next, given a dataset *X* as an *n* × *d* matrix, we will typically interpret *X* as a collection of *n* points {*x_1_, x_2_*, … *x_n_* } in *d*-dimensional space. Finally, for any positive integer *k* ≥ 1, the collection of *top*-*k* nearest neighbors for any point *x*_*i*_, denoted *top* (*k*, *x*_*i*_), is the collection of the *k* points closest to *x*_*i*_ in the sense of the Euclidean norm. The standard kNNG on *X* is defined to be the graph with node set {*x_1_, x_2_*, … *x_n_* } and edge set:

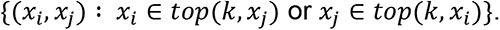

A related construction is the *reciprocal k*-nearest neighbor graph (rkNNG) construction. Here the nodes are given as before, but a stricter convention is followed for the edge set:

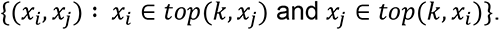

The reciprocal construction takes more information about local densities into account than the standard kNNG, and is occasionally more useful in practice (Qin et al., 2011).

With this setup in place, we are ready to describe our construction. In our setting, datapoints that are close in time also have a tendency to be close in space, as the states change continuously (i.e. without sharp jumps) within an attractor. Keeping this tendency in mind, we carry out the following procedure:

- Construct a standard kNNG
- Remove edges of the form (*x*_*i*_, *x*_*i*+1_) (these will be added back in)
- Remove all non-reciprocal connections, i.e. only retain edges (*x*_*i*_, *x*_j_) if *x*_*i*_ ∈ *top*(*k*, *x*_j_) and *x*_j_ ∈ *top*(*k*, *x*_*i*_)
- Add directed temporal links (*x*_*i*_, *x*_*i*+1_); existing undirected edges (*x*_*i*_, *x*_*j*_) are viewed as double edges (*x*_*i*_, *x*_*j*_), (*x*_*j*_, *x*_*i*_). This final digraph is denoted 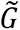.

The initial pruning of temporal neighbors is carried out to help recover directionality in the data (see Figure S5). Other strategies may be possible, but the current method worked sufficiently well for our purposes. The intermediate step of removing non-reciprocal connections is a strategy for disambiguating between points that have differing local densities (Qin et al., 2011). The final addition of the temporal links injects the “arrow of time” back into the graphical representation. Note that this final step is not typical, but is possible in our setting because we work with time series data and know an explicit ordering of our data points. An important feature of the networks that we constructed was that they tended to be *strongly connected* as digraphs, meaning that it was possible to take a directed path between any two vertices. Our construction also accommodates multiple time series or time series with missing frames (frame censoring is common in fMRI time series to remove motion artifacts): the arrow of time is only removed and reintroduced between consecutive time points in the *same* time series or epoch in step 2 and 4 above. When the data constitute a single uninterrupted time series, the digraph will always have one connected component by virtue of the arrows of time that connect every pair of consecutive time points. When the data contain multiple disjoint time series, it is possible for the digraph to have multiple connected components for certain parameter k as the connectivity across time series depends solely on the existence of reciprocal *spatial* neighbors. If it is desirable for the digraph to have a single connected component, one can choose to (a) increase k until reciprocal kNN exists between any pair of time series, or (b) pre-align the time series to increase the spatial proximity between them.

The final step in the construction of the transition network achieves *compression* and follows the notion of *reciprocal clustering of digraphs* introduced by Carlsson et al., (2013). For a fixed parameter *δ* > 0 (specifically *δ* = 2 in this work), we construct an auxiliary undirected graph *U* on *X* with edges 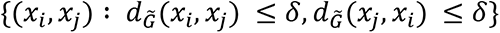. The connected components of *U* partition the vertex set of 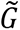 into blocks *B_1_*, *B_2_*, …, *B_m_*. The final compressed transition network *G* is now constructed with vertex set *V(G) B_1_*, *B_2_*, …, *B_m_* and edge set

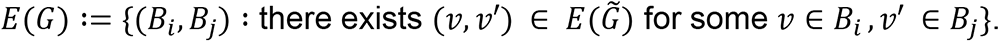

Note that if *v* and *v*^′^belongs to the same connected component of the kNNG 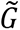, there exists some edge between the partitions of the said connected component. Thus, the number of connected components does not change due to compression.

### 4.5 Gromov-Wasserstein distance between transition networks

Temporal Mapper produces directed graphs where arrows display temporal structure. In general, comparing such graphs directly requires solving a correspondence problem as different graphs may have different numbers of nodes. To solve this correspondence problem, we use the Gromov-Wasserstein (GW) distance (Mémoli, 2007). While originally formulated for metric spaces, the GW formulation was shown to admit a bona fide distance between directed graphs with arbitrary edge weights in (Chowdhury and Mémoli, 2019), and has recently enjoyed significant attention from the machine learning community (Flamary et al., 2021; Peyré et al., 2016; Peyré and Cuturi, 2019; Solomon et al., 2016; Titouan et al., 2019; Xu et al., 2019). In the (di)graph setting, the GW distance allows one to compare the full structure of two (di)graphs without reducing to summary statistics such as degree distributions or centrality.

The GW distance formulation for graphs proceeds as follows. Let *G* = (*V, E*), *H* = (*W*, *F*) be two graphs (possibly directed and/or weighted) on vertex sets *V*, *W* of possibly different sizes. For Temporal Mapper, *E* and *F* are the (asymmetric) geodesic distance matrices–note that these matrices are well-defined because the underlying digraphs are strongly connected. Additionally, let *p*, *q* be two probability distributions on *V*, *W* respectively, denoting the significance of each node. In the Temporal Mapper setting, this is just the number of data points in each node, appropriately normalized, and thus reflects the compression at each node. In matrix-vector notation, we have:

- *E* a |*V*| × |*V*| matrix, *p* a |*V*| × 1 vector such that *p*_*i*_ > 0 for all 1 ≤ *i* ≤ |*V*| and 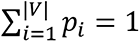
- *F* a |*W*| × |*W*| matrix, *q* a |*W*| × 1 vector such that *q*_*i*_ > 0 for all 1 ≤ *i* ≤ |*W*| and 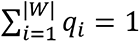

The correspondence between nodes is represented as a joint probability distribution matrix *C* of size |*V*| × |*W*| satisfying nonnegativity and summation constraints *C_ij_* ≥ 0, ∑_*i*,j_ *C_ij_* = 1 as well as marginal constraints such that the rows and columns of *C* correspond to *p* and *q*, respectively. Such a joint distribution is typically referred to as a *coupling matrix*, and the collection of coupling matrices is denoted *C*(*p*, *q*). Intuitively, a coupling describes “how much each node in *V* corresponds to a given node in *W*”.

Finally, the GW distance between the two graphs is a given as the result of the following optimization problem:

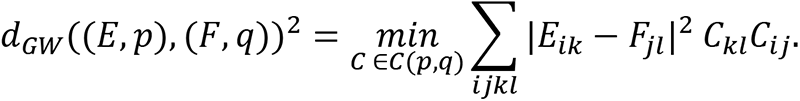

This is a nonconvex quadratic optimization problem (nonconvex because symmetries in the graphs may lead to different correspondences achieving the same minimum, quadratic because the *C* appears twice) and is generally difficult to solve. Intuitively, this distance measures the expected distortion that the edges of *G* would necessarily undergo upon being transformed into the graph *H*. While significant progress has been made in obtaining local minimizers of the underlying optimization through regularization or gradient descent (Flamary et al., 2021; Peyré and Cuturi, 2019), it is in general difficult to assess the quality of these local minimizers except in certain domain areas such as computer vision where the output can be directly inspected in two or three dimensions.

Instead, we opt to solve a lower bound for the GW problem that can be formulated as a linear program with an exact solution. This lower bound, which was referred to as the third lower bound (TLB) in (Mémoli, 2007), arises by decoupling the quadratic optimization problem and instead solving:

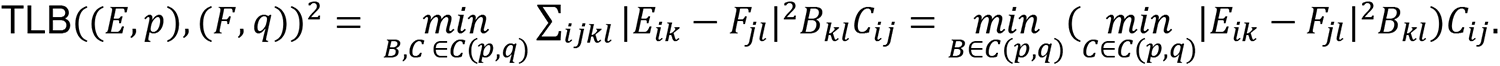

It can be shown (Schmitzer and Schnörr, 2013) that the inner infimization problem has a closed form solution. In other words, the preceding problem amounts to solving the following *linear program*:

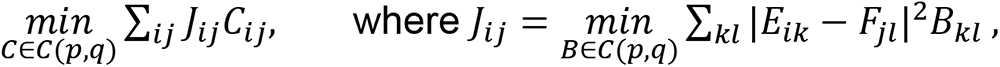

and each *J*_*ij*_ can be individually solved in closed form.

Because there are |*V*| × |*W*| individual entries making up |*J*|, and the entries can all be computed independently, this problem is perfectly suited for GPU computations. Our GPU implementation of the TLB computes each coefficient *J*_*i*j_ in a separate thread block so that all coefficients are computed in parallel. The final linear program can be solved easily using standard linear solvers, e.g. using the network simplex algorithm (Bonneel et al., 2011). For applications in the present work, the GPU-infused version accelerates the original implementation (Chowdhury and Mémoli, 2019) by roughly 200 times.

### 4.6 Cycles of transition networks

Enumerating all cycles in a graph can be computationally expensive (Giscard et al., 2019). However, the transition networks that appear in our setting are small enough that we can enumerate all cycles and carry out further postprocessing in reasonable time. Given a transition network *G* with vertices indexed as {*v*_1_, *v*_2_, …, *v*_*d*_}, we proceed via the following heuristic approach. First we loop over all pairs of vertices (*v*_*i*_, *v*_*j*_) and use Matlab’s native shortest path algorithm (i.e. a breadth-first search of complexity *O*(|*V*| + |*E*|)) to find the shortest paths from *v*_*i*_ to *v*_*j*_ and from *v*_*j*_ to *v*_*i*_, respectively. These paths are then concatenated (avoiding trivial repetition at endpoints) to obtain a cycle. If a cycle has repeated nodes, i.e. is not a simple cycle, then it is discarded. Finally, after the loop terminates, there may be multiple copies of the same cycle with different starting points. For each of these cases, we retain the copy starting at the smallest index and discard the others.

### 4.7 The continuous multitask experiment

In this study, we utilized an fMRI dataset comprising 18 participants collected by (Gonzalez-Castillo et al., 2015) using a continuous multitask paradigm (CMP). We retrieved the data from the XNAT Central public repository (https://central.xnat.org; Project ID: FCStateClassif). Informed consent was obtained from all participants, and the local Institutional Review Board of the National Institute of Mental Health in Bethesda, MD reviewed and approved the CMP data collection. We briefly describe the experiment structure and preprocessing below.

Participants were scanned continuously for 25 minutes and 24 seconds while performing four different cognitive tasks. Each task was presented for two separate 180s blocks, and each task block was preceded by a 12s instruction period. These four tasks were: (1) Rest (R), where participants were instructed to fixate on a crosshair in the center of the screen and let their mind wander; (2) 2-back Working Memory (M), where participants were presented with a continuous sequence of individual geometric shapes and were instructed to press a button whenever the current shape was the same as the shape that appeared two shapes before; (3) Math/arithmetic (A), where participants were presented with simple arithmetic operations, involving three numbers between 1 and 10 and two operands (either addition or subtraction); and (4) Video (V), where participants watched a video of a fish tank from a single point of view with different types of fish swimming into an out of the frame, and were instructed to press a button when a red crosshair appeared on a clown fish and another when it appeared on any other type of fish. For arithmetic, the operations remained on the screen for 4 s and successive trials were separated by a blank screen that appeared for 1 s, yielding a total of 36 operations per each 180s block. For video, the targets appeared for 200 ms with a total of 16 targets during each of the 180s blocks. The order of task blocks was randomized such that the same task did not appear in two consecutive blocks, and the same ordering of tasks was used for all participants. The randomized task order was R-M-V-A-M-R-A-V.

The fMRI data were acquired on a Siemens 7 Tesla MRI scanner equipped with a 32-channel head coil using a whole-brain echo planar imaging (EPI) sequence (repetition time [TR] = 1.5 s, echo time [TE] = 25 ms, and voxel size = 2 mm isotropic). A total of 1017 volumes were acquired while participants performed the continuous multitask paradigm.

Functional and anatomical MR images were preprocessed using the Configurable Pipeline for the Analysis of Connectomes (C-PAC version 0.3.4; https://fcp-indi.github.io/docs/user/index.html). We used the preprocessing utilized in a previous study (Saggar et al., 2018). Briefly, the fMRI data preprocessing steps included ANTS registration into MNI152 space, slice timing correction, motion correction, skull stripping, grand mean scaling, spatial smoothing (FWHM of 4 mm), and temporal band-pass filtering (0.009 Hz < *f* < 0.08Hz). For each ROI, nuisance signal correction was performed by regressing out linear and quadratic trends, physiological noise (white matter and cerebrospinal fluid), motion-related noise (three translational and three rotational head-motion parameters) using the Volterra expansion (Friston et al., 1996) (i.e., six parameters, their temporal derivatives, and each of these values squared), and residual signal unrelated to neural activity extracted using the CompCor algorithm (Behzadi et al., 2007) (i.e., five principal components derived from noise regions in which the time-series data were unlikely to be modulated by neural activity). The resulting data were brought to 3 mm MNI space, and the mean time series was extracted from 375 pre-defined regions-of-interest (ROIs) using the Shine et al. (Shine et al., 2016) atlas. The atlas includes 333 cortical regions from the Gordon et al. (Gordon et al., 2016) atlas, 14 subcortical regions from the Harvard-Oxford subcortical atlas, and 28 cerebellar regions from the SUIT atlas (Diedrichsen et al., 2009). Individual ROIs with zero variance were excluded prior to computing attractor transition networks.

The behavioral data included both responses and reaction times for Working Memory, Math, and Video tasks. Participants were instructed to respond as quickly and accurately as possible with only one response per question. Behavior scores including the percent correct, percent missed, and response times for Working Memory (M), Math (A), and Video (V) tasks were computed for each participant.

## References

Abbott, L.F., Chance, F.S., 2005. Drivers and modulators from push-pull and balanced synaptic input. Prog. Brain Res. 149, 147–155.

Allen, E.A., Damaraju, E., Plis, S.M., Erhardt, E.B., Eichele, T., Calhoun, V.D., 2014. Tracking whole-brain connectivity dynamics in the resting state. Cereb. Cortex 24, 663–676.

Baker, A.P., Brookes, M.J., Rezek, I.A., Smith, S.M., Behrens, T., Probert Smith, P.J., Woolrich, M., 2014. Fast transient networks in spontaneous human brain activity. Elife 3, e01867.

Ban, H., Kalies, W.D., 2006. A Computational Approach to Conley’s Decomposition Theorem. Journal of Computational and Nonlinear Dynamics. https://doi.org/10.1115/1.2338651

Barber, A.D., Lindquist, M.A., DeRosse, P., Karlsgodt, K.H., 2018. Dynamic Functional Connectivity States Reflecting Psychotic-like Experiences. Biological Psychiatry: Cognitive Neuroscience and Neuroimaging 3, 443–453.

Behzadi, Y., Restom, K., Liau, J., Liu, T.T., 2007. A component based noise correction method (CompCor) for BOLD and perfusion based fMRI. Neuroimage 37, 90–101.

Belkin, M., Niyogi, P., 2003. Laplacian Eigenmaps for Dimensionality Reduction and Data Representation. Neural Comput. 15, 1373–1396.

Bonneel, N., van de Panne, M., Paris, S., Heidrich, W., 2011. Displacement interpolation using Lagrangian mass transport, in: Proceedings of the 2011 SIGGRAPH Asia Conference, SA ’11. Association for Computing Machinery, New York, NY, USA, pp. 1–12.

Breakspear, M., 2017. Dynamic models of large-scale brain activity. Nat. Neurosci. 20, 340–352.

Buxton, R.B., Wong, E.C., Frank, L.R., 1998. Dynamics of blood flow and oxygenation changes during brain activation: the balloon model. Magn. Reson. Med. 39, 855– 864.

Buzsaki, G., 2006. Rhythms of the Brain. Oxford University Press.

Cabral, J., Kringelbach, M.L., Deco, G., 2017. Functional connectivity dynamically evolves on multiple time-scales over a static structural connectome: Models and mechanisms. Neuroimage 160, 84–96.

Carlsson, G., 2009. Topology and data. Bull. Am. Math. Soc. 46, 255–308.

Carlsson, G., Mémoli, F., Ribeiro, A., Segarra, S., 2013. Axiomatic construction of hierarchical clustering in asymmetric networks, in: 2013 IEEE International Conference on Acoustics, Speech and Signal Processing. pp. 5219–5223.

Cavanna, F., Vilas, M.G., Palmucci, M., Tagliazucchi, E., 2018. Dynamic functional connectivity and brain metastability during altered states of consciousness. Neuroimage 180, 383–395.

Chazal, F., Michel, B., 2021. An Introduction to Topological Data Analysis: Fundamental and Practical Aspects for Data Scientists. Front Artif Intell 4, 667963.

Chowdhury, S., Mémoli, F., 2019. The Gromov–Wasserstein distance between networks and stable network invariants. Inf Inference 8, 757–787.

Civier, O., Smith, R.E., Yeh, C.-H., Connelly, A., Calamante, F., 2019. Is removal of weak connections necessary for graph-theoretical analysis of dense weighted structural connectomes from diffusion MRI? Neuroimage 194, 68–81.

Cummins, B., Gedeon, T., Harker, S., Mischaikow, K., Mok, K., 2016. Combinatorial Representation of Parameter Space for Switching Networks. SIAM J. Appl. Dyn. Syst. 15, 2176–2212.

Deco, G., Jirsa, V.K., 2012. Ongoing Cortical Activity at Rest: Criticality, Multistability, and Ghost Attractors. Journal of Neuroscience. https://doi.org/10.1523/jneurosci.2523-11.2012

Deco, G., Jirsa, V.K., McIntosh, A.R., 2011. Emerging concepts for the dynamical organization of resting-state activity in the brain. Nat. Rev. Neurosci. 12, 43–56.

Deco, G., Ponce-Alvarez, A., Hagmann, P., Romani, G.L., Mantini, D., Corbetta, M., 2014. How Local Excitation–Inhibition Ratio Impacts the Whole Brain Dynamics. J. Neurosci. 34, 7886–7898.

Deco, G., Ponce-Alvarez, A., Mantini, D., Romani, G.L., Hagmann, P., Corbetta, M., 2013. Resting-state functional connectivity emerges from structurally and dynamically shaped slow linear fluctuations. J. Neurosci. 33, 11239–11252.

Diedrichsen, J., Balsters, J.H., Flavell, J., Cussans, E., Ramnani, N., 2009. A probabilistic MR atlas of the human cerebellum. Neuroimage 46, 39–46.

Díez-Cirarda, M., Strafella, A.P., Kim, J., Peña, J., Ojeda, N., Cabrera-Zubizarreta, A., Ibarretxe-Bilbao, N., 2018. Dynamic functional connectivity in Parkinson’s disease patients with mild cognitive impairment and normal cognition. Neuroimage Clin 17, 847–855.

Du, M., Zhang, L., Li, L., Ji, E., Han, X., Huang, G., Liang, Z., Shi, L., Yang, H., Zhang, Z., 2021. Abnormal transitions of dynamic functional connectivity states in bipolar disorder: A whole-brain resting-state fMRI study. J. Affect. Disord. 289, 7–15.

Edelsbrunner, H., Morozov, D., 2013. Persistent Homology: Theory and Practice. European Congress of Mathematics Kraków, 2 – 7 July, 2012 31–50.

Esfahlani, F.Z., Jo, Y., Faskowitz, J., Byrge, L., Kennedy, D.P., Sporns, O., Betzel, R.F., 2020. High-amplitude cofluctuations in cortical activity drive functional connectivity. Proc. Natl. Acad. Sci. U. S. A. 117, 28393–28401.

Faskowitz, J., Esfahlani, F.Z., Jo, Y., Sporns, O., Betzel, R.F., 2020. Edge-centric functional network representations of human cerebral cortex reveal overlapping system-level architecture. Nat. Neurosci. 23, 1644–1654.

Flamary, R., Courty, N., Gramfort, A., Alaya, M., Laetitia, B.A.C.S., 2021. Pot: Python optimal transport. Journal of Machine Learning Research 22, 1–8.

Fox, M.D., Greicius, M., 2010. Clinical applications of resting state functional connectivity. Front. Syst. Neurosci. 4, 19.

Friston, K.J., Harrison, L., Penny, W., 2003. Dynamic causal modelling. Neuroimage 19, 1273–1302.

Friston, K.J., Mechelli, A., Turner, R., Price, C.J., 2000. Nonlinear responses in fMRI: the Balloon model, Volterra kernels, and other hemodynamics. Neuroimage 12, 466–477.

Friston, K.J., Williams, S., Howard, R., Frackowiak, R.S., Turner, R., 1996. Movement-related effects in fMRI time-series. Magn. Reson. Med. 35, 346–355.

Gameiro, M., Mischaikow, K., Kalies, W., 2004. Topological characterization of spatial-temporal chaos. Phys. Rev. E Stat. Nonlin. Soft Matter Phys. 70, 035203.

Garland, J., Bradley, E., Meiss, J.D., 2016. Exploring the topology of dynamical reconstructions. Physica D 334, 49–59.

Garrity, A.G., Pearlson, G.D., McKiernan, K., Lloyd, D., Kiehl, K.A., Calhoun, V.D., 2007. Aberrant “Default Mode” Functional Connectivity in Schizophrenia. AJP 164, 450–457.

Geniesse, C., Chowdhury, S., Saggar, M., 2022. NeuMapper: A scalable computational framework for multiscale exploration of the brain’s dynamical organization. Network Neuroscience. https://doi.org/10.1162/netn_a_00229

Geniesse, C., Sporns, O., Petri, G., Saggar, M., 2019. Generating dynamical neuroimaging spatiotemporal representations (DyNeuSR) using topological data analysis. Netw Neurosci 3, 763–778.

Giscard, P.-L., Kriege, N., Wilson, R.C., 2019. A General Purpose Algorithm for Counting Simple Cycles and Simple Paths of Any Length. Algorithmica. https://doi.org/10.1007/s00453-019-00552-1

Giusti, C., Pastalkova, E., Curto, C., Itskov, V., 2015. Clique topology reveals intrinsic geometric structure in neural correlations. Proceedings of the National Academy of Sciences. https://doi.org/10.1073/pnas.1506407112

Golos, M., Jirsa, V., Daucé, E., 2015. Multistability in Large Scale Models of Brain Activity. PLoS Comput. Biol. 11, e1004644.

Gonzalez-Castillo, J., Caballero-Gaudes, C., Topolski, N., Handwerker, D.A., Pereira, F., Bandettini, P.A., 2019. Imaging the spontaneous flow of thought: Distinct periods of cognition contribute to dynamic functional connectivity during rest. Neuroimage 202, 116129.

Gonzalez-Castillo, J., Hoy, C.W., Handwerker, D.A., Robinson, M.E., Buchanan, L.C., Saad, Z.S., Bandettini, P.A., 2015. Tracking ongoing cognition in individuals using brief, whole-brain functional connectivity patterns. Proc. Natl. Acad. Sci. U. S. A. 112, 8762–8767.

Gordon, E.M., Laumann, T.O., Adeyemo, B., Huckins, J.F., Kelley, W.M., Petersen, S.E., 2016. Generation and Evaluation of a Cortical Area Parcellation from Resting-State Correlations. Cereb. Cortex 26, 288–303.

Gordon, E.M., Laumann, T.O., Gilmore, A.W., Newbold, D.J., Greene, D.J., Berg, J.J., Ortega, M., Hoyt-Drazen, C., Gratton, C., Sun, H., Hampton, J.M., Coalson, R.S., Nguyen, A.L., McDermott, K.B., Shimony, J.S., Snyder, A.Z., Schlaggar, B.L., Petersen, S.E., Nelson, S.M., Dosenbach, N.U.F., 2017. Precision Functional Mapping of Individual Human Brains. Neuron 95, 791–807.e7.

Hagmann, P., Cammoun, L., Gigandet, X., Meuli, R., Honey, C.J., Wedeen, V.J., Sporns, O., 2008. Mapping the Structural Core of Human Cerebral Cortex. PLoS Biology. https://doi.org/10.1371/journal.pbio.0060159

Hansen, E.C.A., Battaglia, D., Spiegler, A., Deco, G., Jirsa, V.K., 2015. Functional connectivity dynamics: Modeling the switching behavior of the resting state. NeuroImage. https://doi.org/10.1016/j.neuroimage.2014.11.001

Hutchison, R.M., Womelsdorf, T., Allen, E.A., Bandettini, P.A., Calhoun, V.D., Corbetta, M., Della Penna, S., Duyn, J.H., Glover, G.H., Gonzalez-Castillo, J., Handwerker, D.A., Keilholz, S., Kiviniemi, V., Leopold, D.A., de Pasquale, F., Sporns, O., Walter, M., Chang, C., 2013. Dynamic functional connectivity: promise, issues, and interpretations. Neuroimage 80, 360–378.

Kalies, W.D., Mischaikow, K., VanderVorst, R.C.A.M., 2005. An Algorithmic Approach to Chain Recurrence. Found. Comut. Math. 5, 409–449.

Kelso, J.A.S., 2012. Multistability and metastability: understanding dynamic coordination in the brain. Philos. Trans. R. Soc. Lond. B Biol. Sci. 367, 906–918.

Kelso, J.A.S., 1995. Dynamic Patterns: The Self-organization of Brain and Behavior. MIT Press.

Kim, W., Mémoli, F., 2021. Spatiotemporal Persistent Homology for Dynamic Metric Spaces. Discrete Comput. Geom. 66, 831–875.

Laumann, T.O., Snyder, A.Z., Mitra, A., Gordon, E.M., Gratton, C., Adeyemo, B., Gilmore, A.W., Nelson, S.M., Berg, J.J., Greene, D.J., McCarthy, J.E., Tagliazucchi, E., Laufs, H., Schlaggar, B.L., Dosenbach, N.U.F., Petersen, S.E., 2017. On the Stability of BOLD fMRI Correlations. Cereb. Cortex 27, 4719–4732.

Leonardi, N., Van De Ville, D., 2015. On spurious and real fluctuations of dynamic functional connectivity during rest. NeuroImage. https://doi.org/10.1016/j.neuroimage.2014.09.007

Li, J., Zhang, D., Liang, A., Liang, B., Wang, Z., Cai, Y., Gao, M., Gao, Z., Chang, S., Jiao, B., Huang, R., Liu, M., 2017. High transition frequencies of dynamic functional connectivity states in the creative brain. Sci. Rep. 7, 46072.

Lui, S., Wu, Q., Qiu, L., Yang, X., Kuang, W., Chan, R.C.K., Huang, X., Kemp, G.J., Mechelli, A., Gong, Q., 2011. Resting-state functional connectivity in treatment-resistant depression. Am. J. Psychiatry 168, 642–648.

Mandeville, J.B., Marota, J.J., Ayata, C., Zaharchuk, G., Moskowitz, M.A., Rosen, B.R., Weisskoff, R.M., 1999. Evidence of a cerebrovascular postarteriole windkessel with delayed compliance. J. Cereb. Blood Flow Metab. 19, 679–689.

Meer, J.N. van der Breakspear, M., Chang, L.J., Sonkusare, S., Cocchi, L., 2020. Movie viewing elicits rich and reliable brain state dynamics. Nat. Commun. 11, 5004.

Mémoli, F., 2007. On the use of Gromov-Hausdorff distances for shape comparison.

Munch, E., 2013. Applications of persistent homology to time varying systems.

Munn, B. R., Müller, E. J., Wainstein, G. & Shine, J. M., 2021. The ascending arousal system shapes neural dynamics to mediate awareness of cognitive states. Nat Commun 12, 6016.

Myers, A., Munch, E., Khasawneh, F.A., 2019. Persistent homology of complex networks for dynamic state detection. Phys Rev E 100, 022314.

Ou, J., Xie, L., Wang, P., Li, X., Zhu, D., Jiang, R., Wang, Y., Chen, Y., Zhang, J., Liu, T., 2013. Modeling brain functional dynamics via hidden Markov models, in: 2013 6th International IEEE/EMBS Conference on Neural Engineering (NER). pp. 569–572.

Perea, J.A., 2019. Topological Time Series Analysis. Not. Am. Math. Soc. 66, 686–694.

Petri, G., Expert, P., Turkheimer, F., Carhart-Harris, R., Nutt, D., Hellyer, P.J., Vaccarino, F., 2014. Homological scaffolds of brain functional networks. J. R. Soc. Interface 11, 20140873.

Peyré, G., Cuturi, M., 2019. Computational Optimal Transport: With Applications to Data Science. Now Publishers.

Peyré, G., Cuturi, M., Solomon, J., 2016. Gromov-Wasserstein Averaging of Kernel and Distance Matrices, in: Balcan, M.F., Weinberger, K.Q. (Eds.), Proceedings of The 33rd International Conference on Machine Learning, Proceedings of Machine Learning Research. PMLR, New York, New York, USA, pp. 2664–2672.

Poincaré, H., 1967. New Methods of Celestial Mechanics. National Aeronautics and Space Administration.

Preti, M.G., Bolton, T.A., Van De Ville, D., 2017. The dynamic functional connectome: State-of-the-art and perspectives. Neuroimage 160, 41–54.

Qi, C.R., Su, H., Mo, K., Guibas, L.J., 2017. Pointnet: Deep learning on point sets for 3d classification and segmentation, in: Proceedings of the IEEE Conference on Computer Vision and Pattern Recognition. pp. 652–660.

Qin, D., Gammeter, S., Bossard, L., Quack, T., van Gool, L., 2011. Hello neighbor: Accurate object retrieval with k-reciprocal nearest neighbors, in: CVPR 2011. pp. 777–784.

Quinn, A.J., Vidaurre, D., Abeysuriya, R., Becker, R., Nobre, A.C., Woolrich, M.W., 2018. Task-Evoked Dynamic Network Analysis Through Hidden Markov Modeling. Front. Neurosci. 12, 603.

Rabany, L., Brocke, S., Calhoun, V.D., Pittman, B., Corbera, S., Wexler, B.E., Bell, M.D., Pelphrey, K., Pearlson, G.D., Assaf, M., 2019. Dynamic functional connectivity in schizophrenia and autism spectrum disorder: Convergence, divergence and classification. Neuroimage Clin 24, 101966.

Rabiner, L.R., 1989. A tutorial on hidden Markov models and selected applications in speech recognition. Proc. IEEE 77, 257–286.

Rezek, I., Roberts, S., 2005. Ensemble Hidden Markov Models with Extended Observation Densities for Biosignal Analysis, in: Husmeier, D., Dybowski, R., Roberts, S. (Eds.), Probabilistic Modeling in Bioinformatics and Medical Informatics. Springer London, London, pp. 419–450.

Saggar, M., Shine, J.M., Liégeois, R., Dosenbach, N.U.F., Fair, D., 2021. Precision dynamical mapping using topological data analysis reveals a unique hub-like transition state at rest. bioRxiv. https://doi.org/10.1101/2021.08.05.455149

Saggar, M., Sporns, O., Gonzalez-Castillo, J., Bandettini, P.A., Carlsson, G., Glover, G., Reiss, A.L., 2018. Towards a new approach to reveal dynamical organization of the brain using topological data analysis. Nat. Commun. 9, 1399.

Saggar, M., Uddin, L.Q., 2019. Pushing the boundaries of psychiatric neuroimaging to ground diagnosis in biology. eNeuro 6, ENEURO.0384–19.2019.

Schmitzer, B., Schnörr, C., 2013. Modelling Convex Shape Priors and Matching Based on the Gromov-Wasserstein Distance. J. Math. Imaging Vis. 46, 143–159.

Shine, J.M., Bissett, P.G., Bell, P.T., Koyejo, O., Balsters, J.H., Gorgolewski, K.J., Moodie, C.A., Poldrack, R.A., 2016. The Dynamics of Functional Brain Networks: Integrated Network States during Cognitive Task Performance. Neuron 92, 544– 554.

Singh, G., Mémoli, F., Carlsson, G.E., Others, 2007. Topological methods for the analysis of high dimensional data sets and 3d object recognition. PBG@ Eurographics 2.

Smith, S.M., Beckmann, C.F., Andersson, J., Auerbach, E.J., Bijsterbosch, J., Douaud, G., Duff, E., Feinberg, D.A., Griffanti, L., Harms, M.P., Kelly, M., Laumann, T., Miller, K.L., Moeller, S., Petersen, S., Power, J., Salimi-Khorshidi, G., Snyder, A.Z., Vu, A.T., Woolrich, M.W., Xu, J., Yacoub, E., Uğurbil, K., Van Essen, D.C., Glasser, M.F., WU-Minn HCP Consortium, 2013. Resting-state fMRI in the Human Connectome Project. Neuroimage 80, 144–168.

Solomon, J., Peyré, G., Kim, V.G., Sra, S., 2016. Entropic metric alignment for correspondence problems. ACM Trans. Graph. 35, 1–13.

Taghia, J., Cai, W., Ryali, S., Kochalka, J., Nicholas, J., Chen, T., Menon, V., 2018. Uncovering hidden brain state dynamics that regulate performance and decision-making during cognition. Nat. Commun. 9, 2505.

Tang, Y.-Y., Rothbart, M.K., Posner, M.I., 2012. Neural correlates of establishing, maintaining, and switching brain states. Trends Cogn. Sci. 16, 330–337.

Tenenbaum, J.B., de Silva, V., Langford, J.C., 2000. A Global Geometric Framework for Nonlinear Dimensionality Reduction. Science. https://doi.org/10.1126/science.290.5500.2319

Titouan, V., Courty, N., Tavenard, R., Laetitia, C., Flamary, R., 2019. Optimal Transport for structured data with application on graphs, in: Chaudhuri, K., Salakhutdinov, R. (Eds.), Proceedings of the 36th International Conference on Machine Learning, Proceedings of Machine Learning Research. PMLR, pp. 6275–6284.

Tognoli, E., Kelso, J.A.S., 2014. The metastable brain. Neuron 81, 35–48.

Topaz, C.M., Ziegelmeier, L., Halverson, T., 2015. Topological data analysis of biological aggregation models. PLoS One 10, e0126383.

Tymochko, S., Munch, E., Khasawneh, F.A., 2020. Using Zigzag Persistent Homology to Detect Hopf Bifurcations in Dynamical Systems. Algorithms 13, 278.

Ulmer, M., Ziegelmeier, L., Topaz, C.M., 2019. A topological approach to selecting models of biological experiments. PLoS One 14, e0213679.

Umeyama, S., 1988. An eigendecomposition approach to weighted graph matching problems. IEEE Trans. Pattern Anal. Mach. Intell. 10, 695–703.

van den Heuvel, M.P., Hulshoff Pol, H.E., 2010. Exploring the brain network: A review on resting-state fMRI functional connectivity. Eur. Neuropsychopharmacol. 20, 519– 534.

Van Der Maaten, L., Postma, E., Van den Herik, J., 2009. Dimensionality reduction: a comparative. J. Mach. Learn. Res.

Van Essen, D.C., Smith, S.M., Barch, D.M., Behrens, T.E.J., Yacoub, E., Ugurbil, K., WU-Minn HCP Consortium, 2013. The WU-Minn Human Connectome Project: an overview. Neuroimage 80, 62–79.

Vidaurre, D., Smith, S.M., Woolrich, M.W., 2017. Brain network dynamics are hierarchically organized in time. Proc. Natl. Acad. Sci. U. S. A. 114, 12827–12832.

Wang, X.-J., 2002. Probabilistic decision making by slow reverberation in cortical circuits. Neuron 36, 955–968.

Wilson, H. R. & Cowan, J. D., 1972. Excitatory and Inhibitory Interactions in Localized Populations of Model Neurons. Biophys J 12, 1–24.

Wilson, H. R. & Cowan, J. D., 1973. A mathematical theory of the functional dynamics of cortical and thalamic nervous tissue. Kybernetik 13, 55–80.

Wong, K.-F., Wang, X.-J., 2006. A recurrent network mechanism of time integration in perceptual decisions. J. Neurosci. 26, 1314–1328.

Xu, H., Luo, D., Carin, L., 2019. Scalable Gromov-Wasserstein learning for graph partitioning and matching. Adv. Neural Inf. Process. Syst.

Zalesky, A., Fornito, A., Cocchi, L., Gollo, L.L., Breakspear, M., 2014. Time-resolved resting-state brain networks. Proc. Natl. Acad. Sci. U. S. A. 111, 10341–10346.

Zamani Esfahlani, F., Jo, Y., Faskowitz, J., Byrge, L., Kennedy, D.P., Sporns, O., Betzel, R.F., 2020. High-amplitude cofluctuations in cortical activity drive functional connectivity. Proc. Natl. Acad. Sci. U. S. A. 117, 28393–28401.

Zaslavskiy, M., Bach, F., Vert, J.-P., 2009. A path following algorithm for the graph matching problem. IEEE Trans. Pattern Anal. Mach. Intell. 31, 2227–2242.

Zhang, M., Kalies, W.D., Kelso, J.A.S., Tognoli, E., 2020. Topological portraits of multiscale coordination dynamics. J. Neurosci. Methods 339, 108672.

Zhang, M., Sun, Y., Saggar, M., 2022. Cross-attractor repertoire provides new perspective on structure-function relationship in the brain. https://doi.org/10.1101/2020.05.14.097196

