## Supplementary Materials for "Temporal Mapper: transition networks in simulated and real neural dynamics"

### 1 Supplementary Materials

#### S1 Region indices and names

| abbreviation | region name | right index | left index |
|---|---|---|---|
| ENT | Entorhinal cortex | 1 | 66 |
| PARH | Parahippocampal cortex | 2 | 65 |
| TP | Temporal pole | 3 | 64 |
| FP | Frontal pole | 4 | 63 |
| FUS | Fusiform gyrus | 5 | 62 |
| TT | Transverse temporal cortex | 6 | 61 |
| LOCC | Lateral occipital cortex | 7 | 60 |
| SP | Superior parietal cortex | 8 | 59 |
| IT | Inferior temporal cortex | 9 | 58 |
| IP | Inferior parietal cortex | 10 | 57 |
| SMAR | Supramarginal gyrus | 11 | 56 |
| BSTS | Bank of the superior temporal sulcus | 12 | 55 |
| MT | Middle temporal cortex | 13 | 54 |
| ST | Superior temporal cortex | 14 | 53 |
| PSTC | Postcentral gyrus | 15 | 52 |
| PREC | Precentral gyrus | 16 | 51 |
| CMF | Caudal middle frontal cortex | 17 | 50 |
| POPE | Pars opercularis | 18 | 49 |
| PTRI | Pars triangularis | 19 | 48 |
| RMF | Rostral middle frontal cortex | 20 | 47 |
| PORB | Pars orbitalis | 21 | 46 |
| LOF | Lateral orbitofrontal cortex | 22 | 45 |
| CAC | Caudal anterior cingulate cortex | 23 | 44 |
| RAC | Rostral anterior cingulate cortex | 24 | 43 |
| SF | Superior frontal cortex | 25 | 42 |
| MOF | Medial orbitofrontal cortex | 26 | 41 |
| LING | Lingual gyrus | 27 | 40 |
| PCAL | Pericalcarine cortex | 28 | 39 |
| CUN | Cuneus | 29 | 38 |
| PARC | Paracentral lobule | 30 | 37 |
| ISTC | Isthmus of the cingulate cortex | 31 | 36 |
| PCUN | Precuneus | 32 | 35 |
| PC | Posterior cingulate cortex | 33 | 34 |

**Table S1. Names and indices of brain regions**. As in [1], the brain is parcellated into 66 regions as shown in Figure 2b. Here is a list of the specific region names corresponding to each region index. Region 1-33 are on the right hemisphere, and region 66-34 are the homologous regions on the the left hemisphere.

#### S2 Additional information regarding the construction from simulated data

To show that the reconstructed transition networks reveal non-stationary information in the original time series, we construct transition networks from phase-randomized time series (1000 trials) using the same construction parameters as those of Figure 3e-h. Comparing the reconstruction from phase-randomized data to the ground truth (Figure 2f,g) yields null distributions shown in Figure S1.

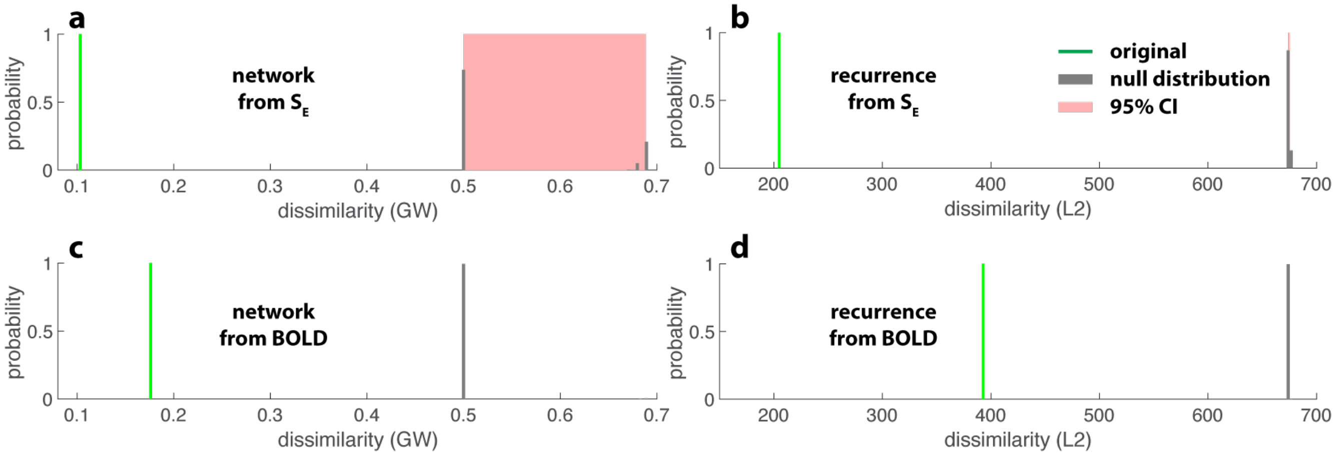


**Figure S1. Differentiation between reconstruction from the original data versus phase-randomized data.** Green lines indicate the reconstruction errors reproduced from Figure 3e-h. Gray bars indicate the null distributions computed from phase-randomized time series (1000 trials) and their corresponding confidence intervals.

Figure S2 shows how the choice of neighborhood size (k) and the contraction distance ($\delta$) influences the reconstruction error for the simulated neural dynamics (a,b) and derived BOLD signals (c,d). For a low level of compression (small $\delta$), the reconstruction error primarily depends on $k$. Over-compression increases the error. For a wide range of parameters, the reconstruction error remains low (near-black regions), indicating the stability of reconstruction.

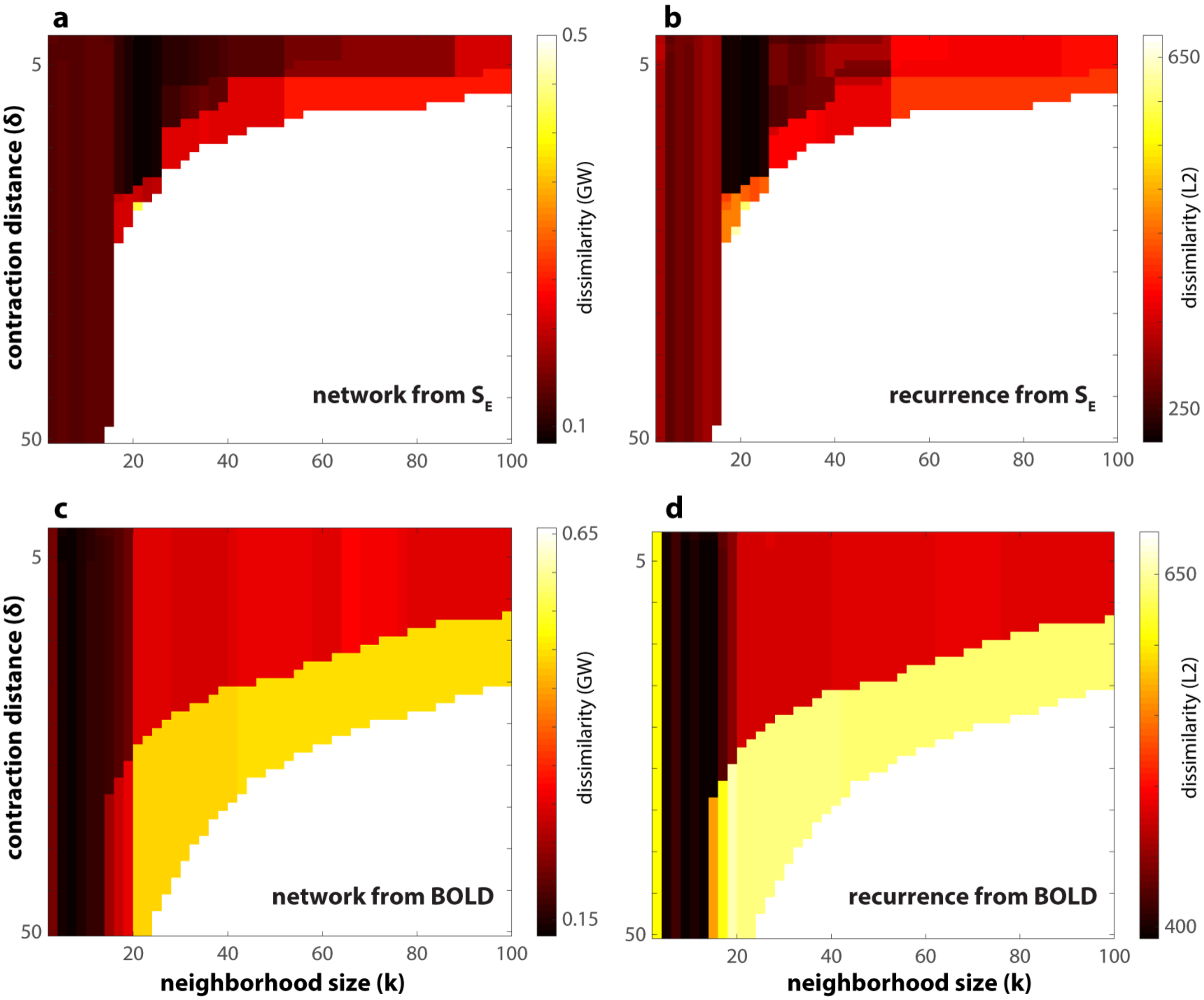


**Figure S2. Parameter-dependency of reconstruction error.** (a-d) extends Figure 3e-h to show how the reconstruction error depends on the construction parameters k—the neighborhood size in the construction of the neighborhood graph (e.g. Figure 3b) and $\delta$—the distance within which the nodes in the neighborhood graph are contracted.

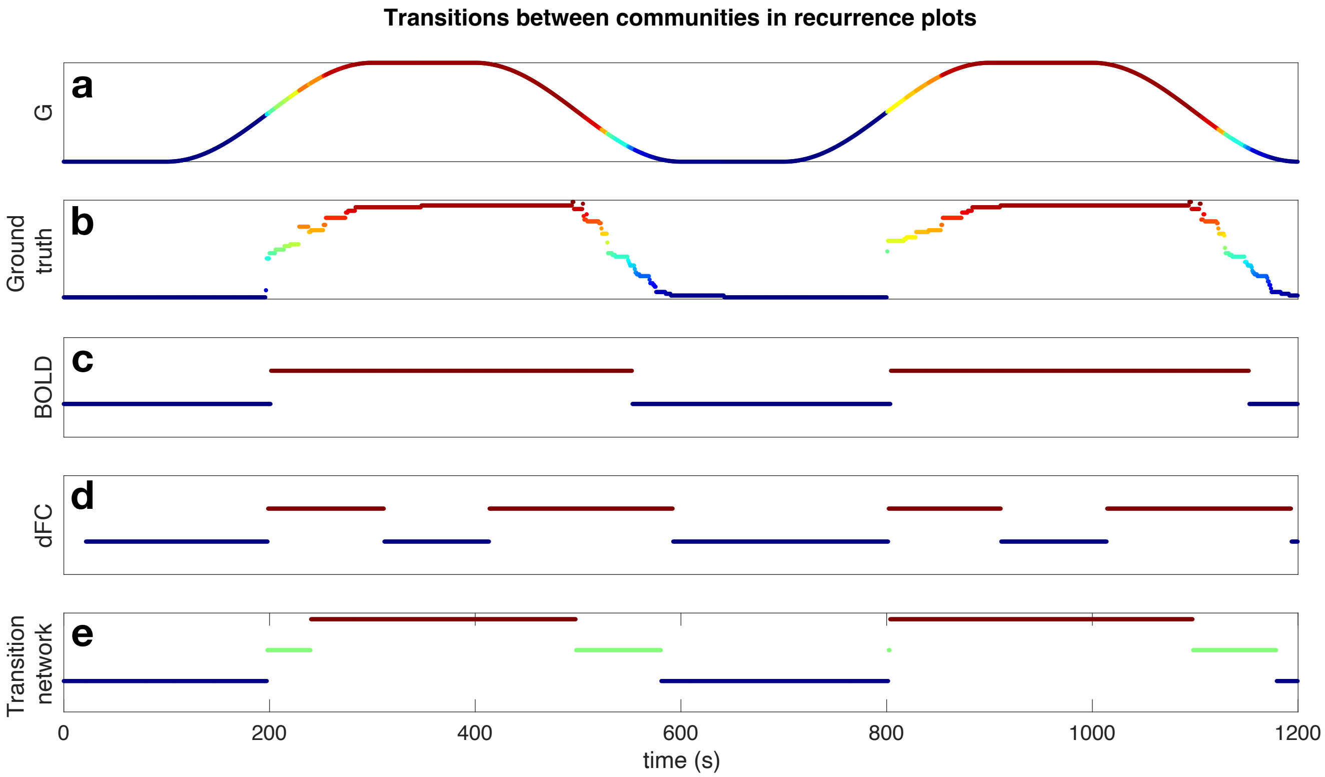


**Figure S3. Performance of transition networks in detecting states.** Following up on Figure 4, we consider an alternative method for characterizing transitions between states based on recurrence plots. Panels (a), (b) show the dynamics of the control parameter $G$ and the ground truth attractor labels after applying sufficient downsampling to retain a visually manageable amount of detail. In this case, there are 55 unique attractors in total. To obtain panels (c-e), we applied the following method to the recurrence plots associated to BOLD, dFC (computed using 30 time frames), and the attractor transition network: first we normalized values to be in $[0,1]$, then applied a transformation $x \mapsto 1-x$ to convert the dissimilarity into a similarity, and then applied the Louvain community detection algorithm [2] to the resulting matrix treated as a weighted graph adjacency matrix. This method detected two states for each of the BOLD and dFC recurrence matrices, and an additional transitory state for the transition network recurrence plot. Panel (c) shows that the two states with longest occupancy were reliably recovered by BOLD. Panel (d) appears to show sensitivity to transitions of $G$. Further computations with different dFC window sizes are illustrated in Figure S4. Finally, panel (e) shows the unsupervised recovery of states via our transition networks. Here we see a third, transitory state bridging the two that are occupied for the longest time. This additional state better captures the timespan during which transitions take place between attractors.

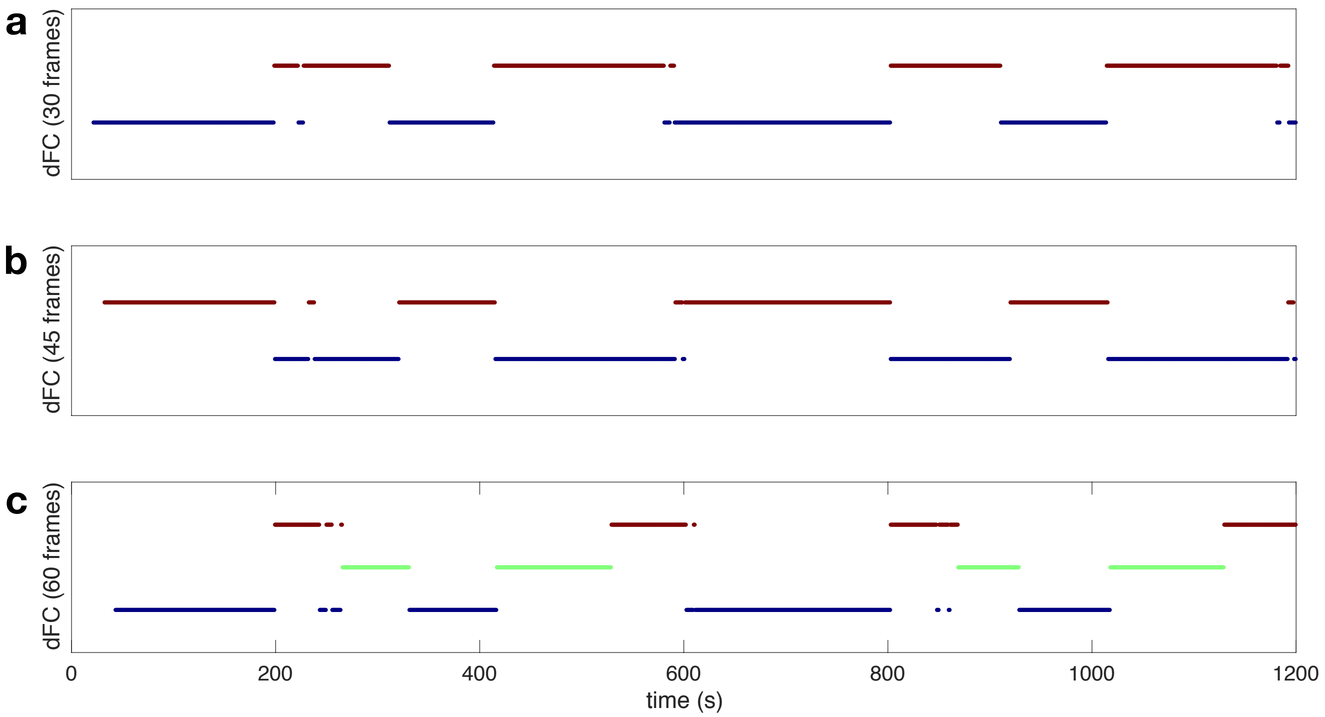


**Figure S4. Parameter perturbation for dFC in detecting states.** Following Figure 4j, we additionally performed community-based state detection on recurrence plots of dFC computed using 45 and 60 time frames (shown in (b) and (c)). While we see that recovery of additional states is possible, it nonetheless appears that dFC-based methods may be better suited for detecting transitions between states rather than stable occupancy of particular states.

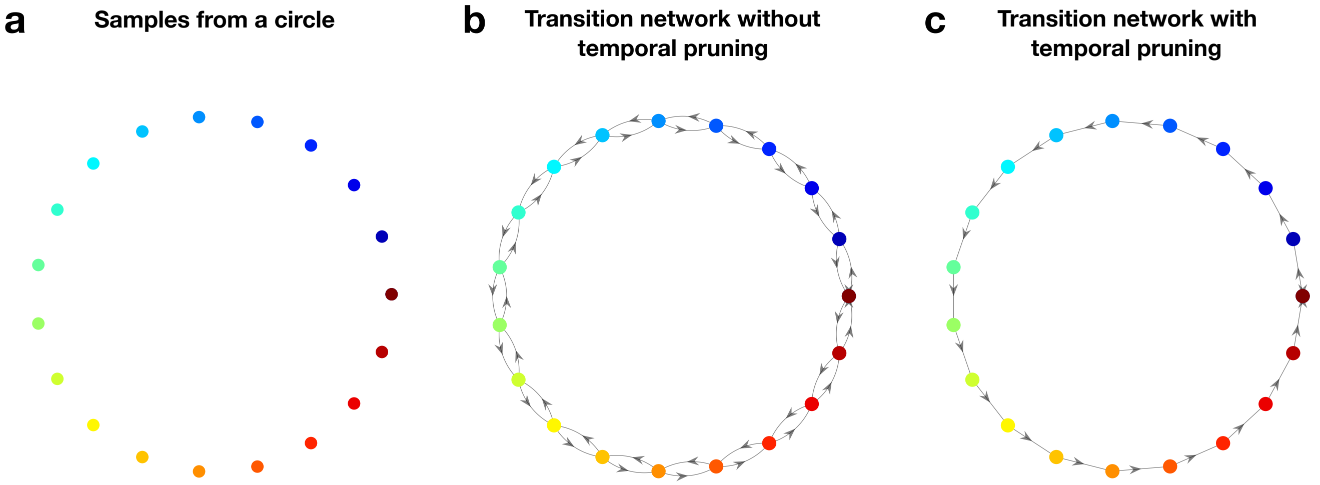


**Figure S5. Effect of temporal pruning in constructing transition networks.** When constructing transition networks, we initially prune temporal neighbors $(x_i, x_{i+1})$ before pruning non-reciprocal connections. To see how this aids in capturing directionality information, consider panel (a), where points are sampled from a circle with sampling order indicated by the color map. Without temporal pruning, the constructed transition network captures spatial information, but not the temporal order in which points were sampled (panel (b)). With temporal pruning, this temporal ordering is recovered in the directionality of the arrows (panel (c)).

#### S3 Additional information regarding the construction from human fMRI data

To examine how the absolute source-sink difference (absolute value of black curve in Figure 5h) differs in different parts of each block and in different tasks. We calculate the average absolute difference within the entry, exit, and middle segments of each block. The absolute difference values are further averaged over all entry and exit segments of each task for each subject; similarly, the middle segments are averaged within each task for each subject. A 2-way ANOVA is performed on the resulted absolute difference values with the type of task and the location of the segment as independent variables. There are significant main effects of both the type of task (F(3,136)=20.85, p<$10^{-10}$) and the local of the segment (F(1,136)=20.81, p=$1.12 \times 10^{-5}$). The post-hoc comparisons are shown in Figure S7 (Tukey-HSD corrected). The interaction between the two independent variables are not statistically significant (F(3,136)=0.24, p=0.869).

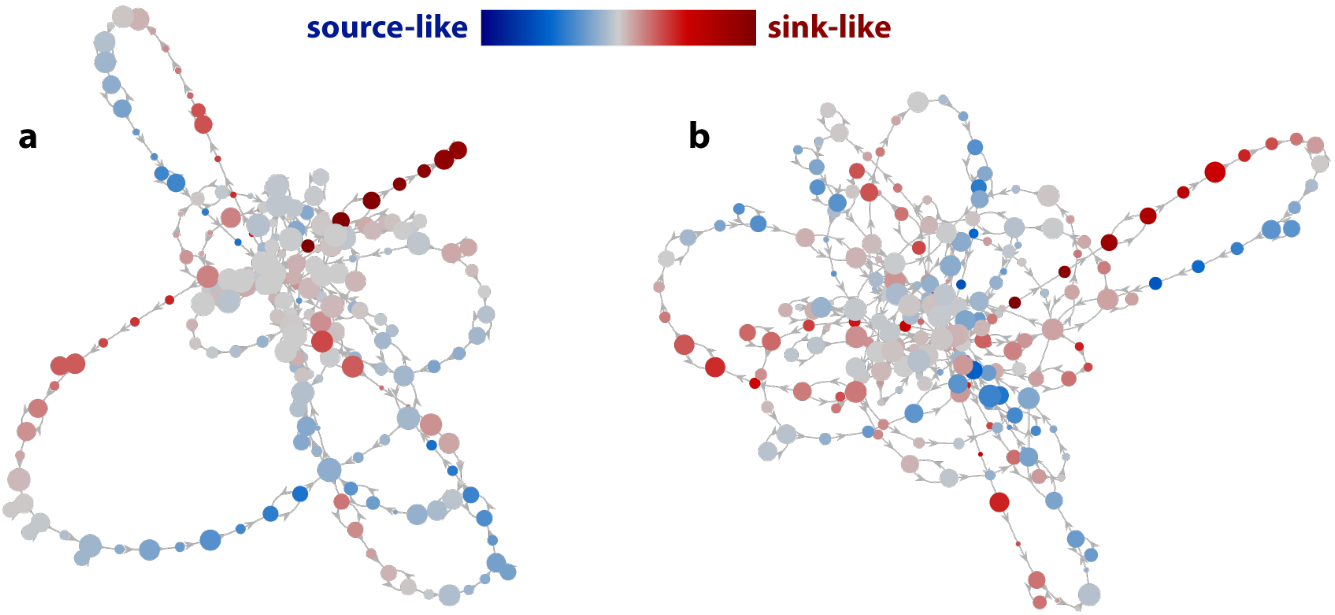


**Figure S6. Examples of source-sink difference of different nodes.** Attractor transition networks of subject 17 (a) and 12 (b), shown in Figure 5, are here recolored by the source-sink difference of each node. Blue indicates negative source-sink difference, or a more source-like node. Red indicates positive source-sink difference, or a more sink-like node. Gray nodes are neutral.

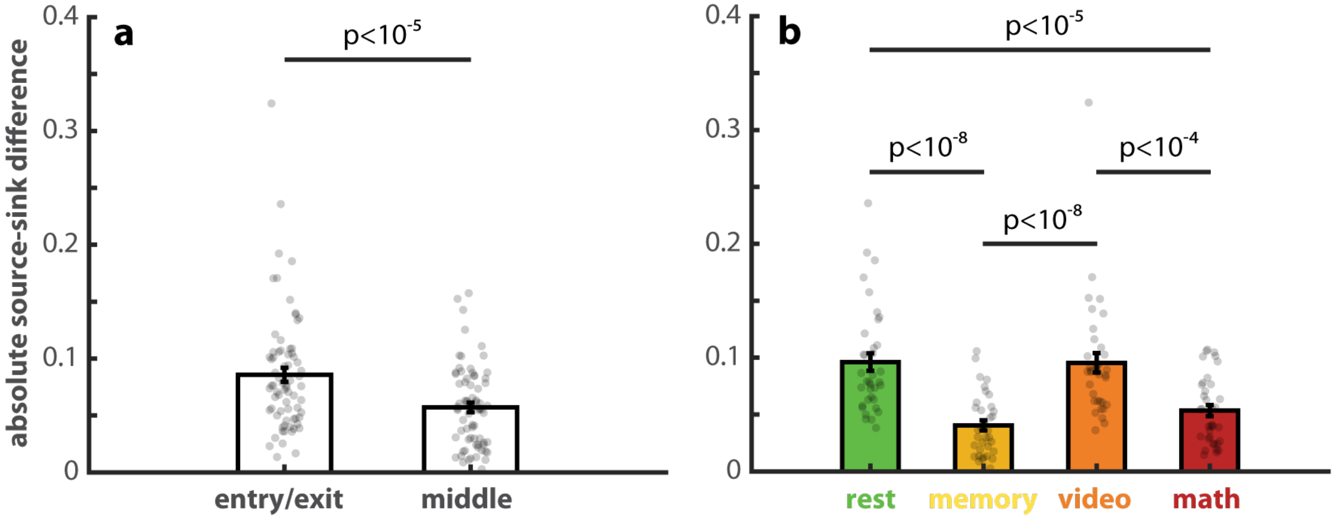


**Figure S7.** **Absolute source-sink difference varies within blocks and across tasks.** We define the first 20 TRs (30s) of each block as the entry and the last 20 TRs as the exit. Between the entry and the exit, there is a middle segment of 80 TRs (120s). (a) shows that the average absolute source-sink difference in the entry and exit is significantly greater than that of the middle segment. Each dot in (a) represent the entry/exit or middle segments of one task and one subject. (b) shows that rest and the video task exhibit a greater absolute source-sink difference than the memory and math tasks. The dots in (b) represent the same information as in (a) but regrouped by tasks.

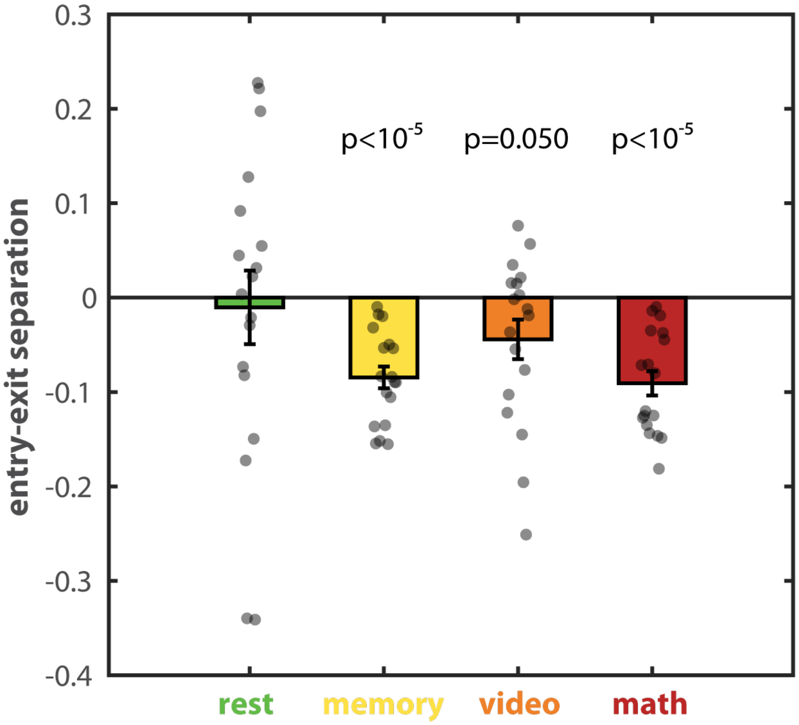


**Figure S8. Entry-exit separation primarily occurs in memory and math tasks.** In Figure 5h, one can observe that in blocks of memory (yellow) and math (red) tasks, brain dynamics enter each block from source-like nodes (negative curve) and exit the block from sink like nodes (positive curve)—an entry-exit separation. In contrast, blocks of rest and video tasks do not consistently exhibit this feature. This figure provides the statistical confirmation of this observation. Quantitatively, we compute the entry-exit separation as the average source-sink difference in the first 20 TRs (30s) minus that of the last 20 TRs of each block; the results are furthered averaged within each subject and each task. A more negative score indicates that the entry (exit) to the block is more source (sink) like. Remarkably, the entry-exit separation is consistently negative for the memory and math tasks across all subjects (1 black = 1 subject). The entry-exit separation is statistically significant in the memory and math tasks, marginally significant for the video task, and non-significant for rest ($p=0.79$).

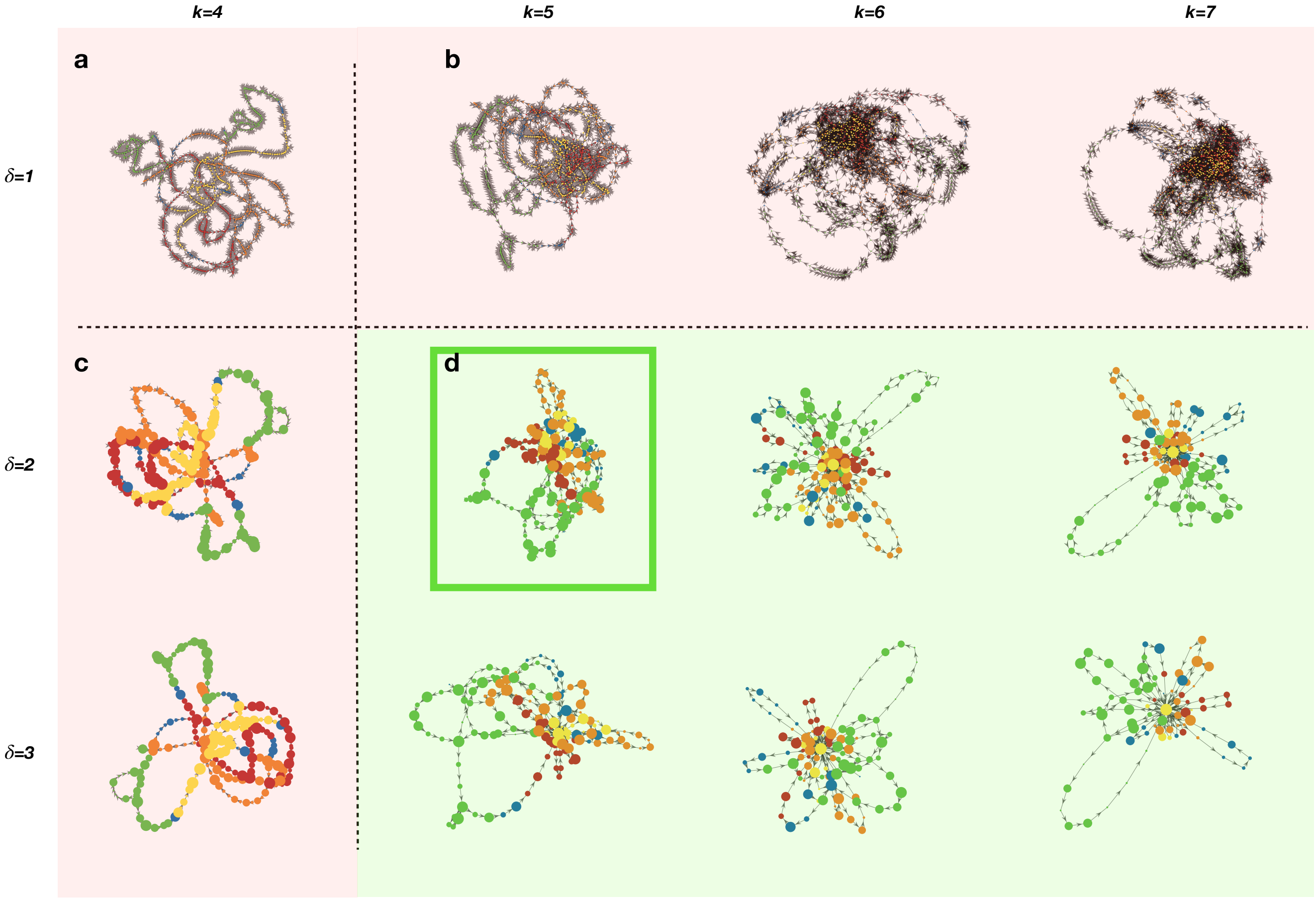


**Figure S9. Parameter perturbation for construction of transition networks.** To determine parameters for transition networks constructed from the human fMRI dataset, we utilized a two-parameter elbow method. For an exemplar subject (Subject 1 shown), we first observed that $\delta = 2$ was a critical parameter at which graph compression became significant (shown in the column containing panels (a), (c)). We also observed that $k = 5$ was a critical parameter at which a substantial core appeared in the graph, which in turn disambiguated between high density and low density regions of the data space (shown in the row containing panels (a), (b)). We thus chose $\delta = 2$ and $k = 5$ as our parameters when reporting results. See also Figure S10 for further validation of these parameter choices.

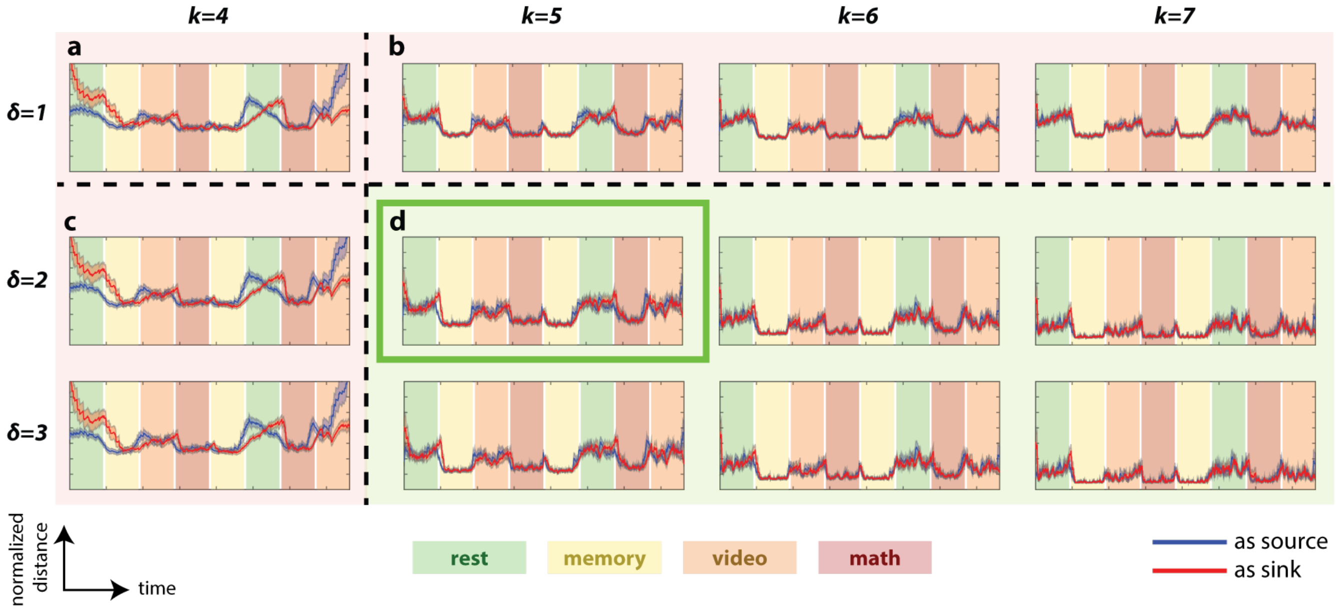


**Figure S10. Effect of parameter perturbation on group level statistics derived from transition networks.** In addition to the visually observable differences at critical parameters, we also observed group-level differences in the ability of transition networks to determine task transitions as a function of parameters. Here, for different $k, \delta$ parameters, we follow Figure 5 and plot the normalized average geodesic distance from: (1) each TR as a source to all other TRs (blue), and (2) to each TR as a sink from all other TRs (red). The normalization is carried out using the maximal distance for each subject. The horizontal axis shows time (in seconds) in the range $[0, 1525]$, and the vertical axis spans the range $[0, 0.7]$. Solid lines show the averages across subjects; shaded areas show the corresponding standard errors. Large changes in the plotted values correspond to task transitions. We observe that for choices of $k$ below the critical value of $k = 5$, the lack of a core (see Figure S9) affects the ability of the transition networks to recover task transitions. These transitions are more stable to choices of $\delta$, but due to the lack of compression in the case $\delta = 1$ (see Figure S9), these plotted statistics require much greater runtime to compute for $\delta = 1$ compared to $\delta \geq 2$. Hence for both scalability and visualization purposes, we consider $\delta = 2$ to be a critical threshold.

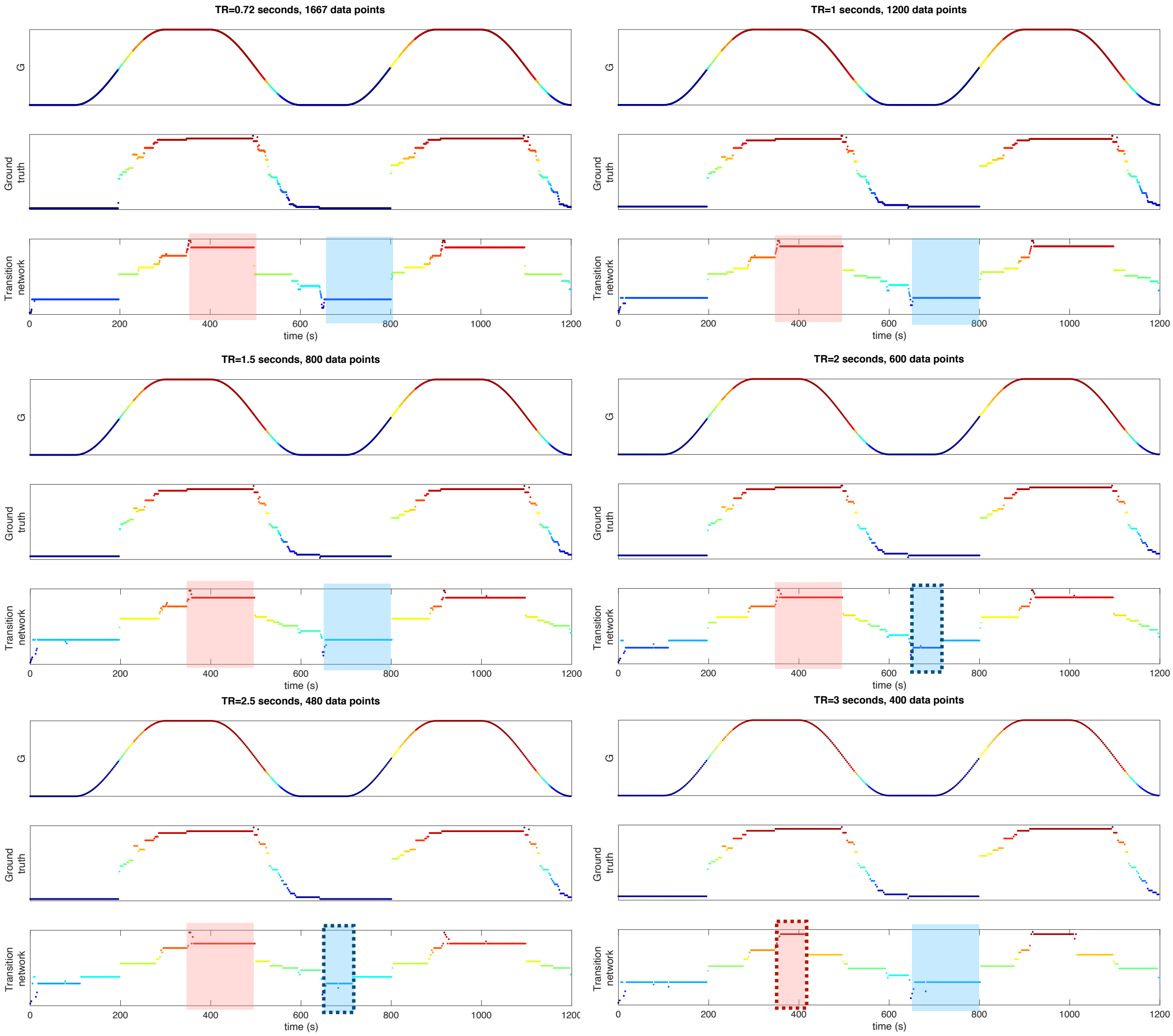


**Figure S11. Resiliency under data ablation.** To better understand the data needs for Temporal Mapper, we repeated the computations presented in Figure 4 after reducing the number of data points via subsampling. The subplots correspond to the variety of TRs typically used in fMRI experiments. We find that the reconstructed transition networks continue to track state dynamics as we increase TR to 3 seconds, albeit with performance degradation (evidenced by shorter state durations, see dotted boxes for some examples) starting at TR>=2s. In particular, Temporal Mapper is able to track state transitions and state durations reasonably well for TR=1.5s, which is also the TR for the real human fMRI dataset we have used in this work.

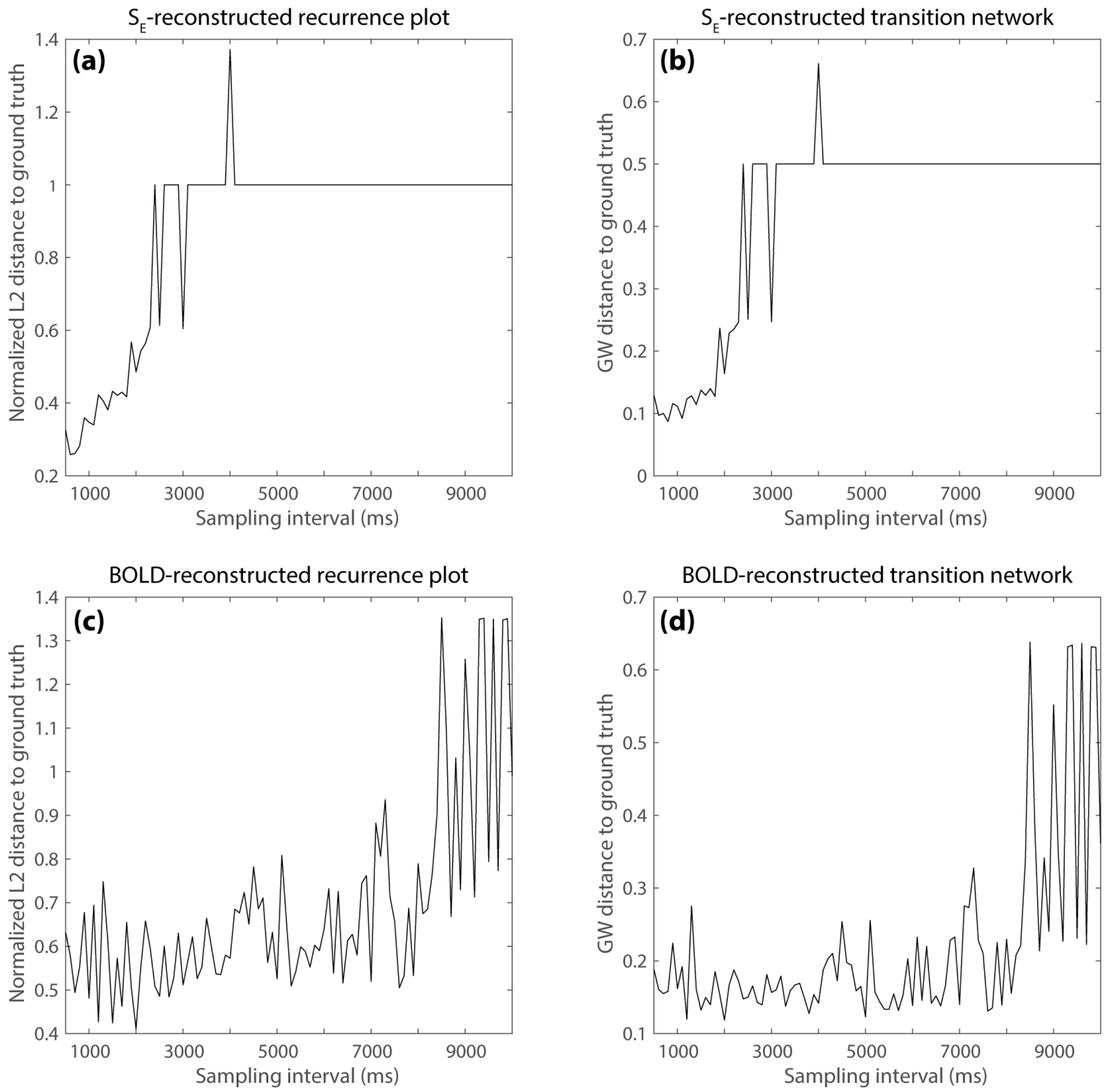


**Figure S12. Reconstruction error by sampling interval.** Quantitative measures of reconstruction errors are shown here in terms of the normalized L2 distance (matrix L2 norm = 1) between the reconstructed recurrence plot and the ground-truth recurrence plot (a and c for reconstructions from $S_E$ and BOLD, respectively) and the Gromov-Wasserstein (GW) distance between the reconstructed attractor transition network and the ground-truth transition network (b and d for reconstructions from $S_E$ and BOLD respectively). Overall, the reconstruction errors first gradually increase with increasing sampling interval (i.e., decreasing sampling rate) until reaching a critical value where the errors jump to the maximum. This critical value is around 2.5 s for $S_E$ and 8 s for BOLD. Interestingly, although the reconstruction error from neural activities $S_E$ is lower than that of BOLD given a short sampling interval (<2 s), BOLD is more robust against down sampling. This may be due to the low-pass filtering effect of the hemodynamic response, where high-frequency fluctuations preserved in $S_E$ are more sensitive to down-sampling.

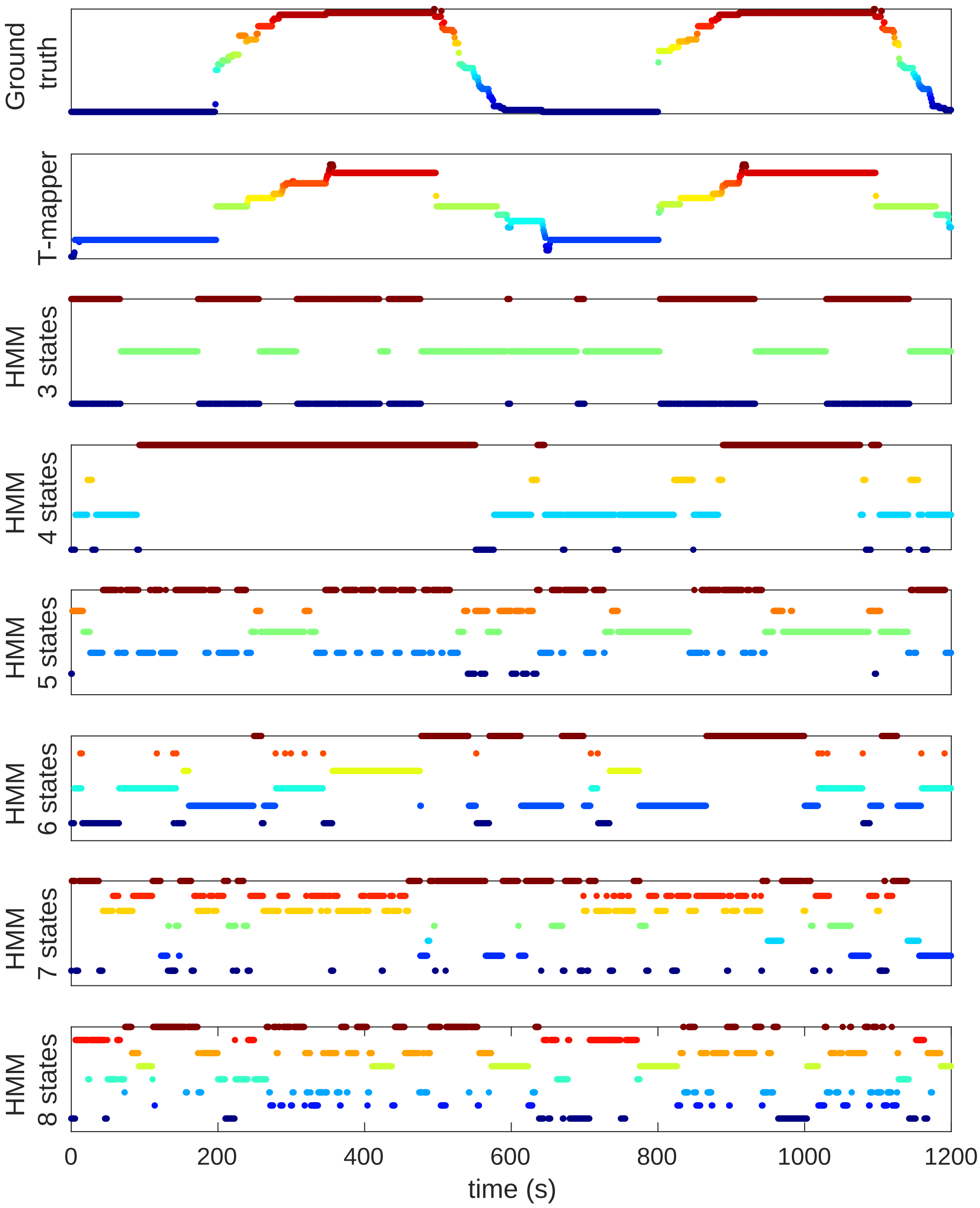


**Figure S13. Comparison with Hidden Markov Models.** Here we repeat the computation of Figure 4 after training and sampling from HMMs with varying numbers of hidden states. Crucially, the Markovian property of a standard HMM implies that the duration for which the modeled process remains at a particular state is modeled by an exponential distribution [3], and consequently, a standard HMM is unable to track the underlying state dynamics shown in Figure 4. Although there are variants of HMMs that explicitly model the time spent in each state–often referred to as Hidden Semi-Markov Models–comparison to these variants is beyond the scope of this work. The HMMs presented here were trained using Matlab's `hmmtrain`. To train, we first z-scored our simulated dataset (following [4]), and then used a k-means method with k=3n—where n is a user-defined number of states—to discretize our simulated dataset comprising $L = 1667$ observations into a set of $k$ emission symbols. Next, we sampled 100 sequences of length $l < L$, where $l$ was uniformly sampled from $[200, 400]$ from the simulated dataset. We used these samples to train our HMM to convergence. Prior to training, we initialized the $n \times n$ state transition matrix and $n \times k$ emission probability matrix with uniform random distributions. After training, we sampled $L$ observations from the HMM. We repeated this procedure for $n = 3, 4, 5, 6, 7, 8$. Note that relative to the HMMs (bottom six), Temporal Mapper (second from top) is much better able to model the state durations of the ground truth transition network (top).

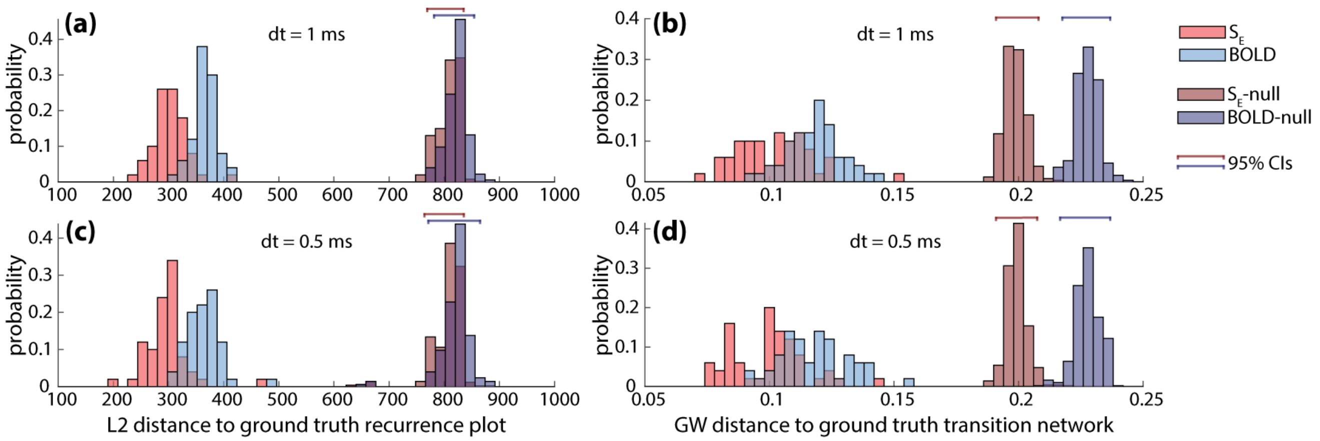


**Figure S14. The effect of noise and the integration time step.** 100 realizations of the model with noise amplitude $\sigma = 0.01$ were simulated using stochastic Heun's method to show the effect of noise and the integration time step (50 for $\mathrm{d}t = 1$ ms and 50 for $\mathrm{d}t = 0.5$ ms). We found no significant difference ($p > 0.25$) between the reconstruction errors for simulations with the integration time step $\mathrm{d}t = 1$ ms and $\mathrm{d}t = 0.5$ ms for either the neural activities $S_E$ (light red bars) or BOLD signal (light blue bars) using different measures (a and c for L2 distance; b and d for Gromov-Wasserstein distance). The distributions of reconstruction errors given noise are far from the 95% confidence intervals of the null distributions (i.e., reconstruction errors for temporally permuted data; dark bars). The reconstruction errors for $S_E$ are significantly lower than those of BOLD ($p < 10^{-8}$).

#### S4 References


1. Deco, G. *et al.* Resting-state functional connectivity emerges from structurally and dynamically shaped slow linear fluctuations. *Journal of Neuroscience* **33,** 11239–11252 (2013).
2. Blondel, V. D., Guillaume, J.-L., Lambiotte, R. & Lefebvre, E. Fast unfolding of communities in large networks. *Journal of statistical mechanics: theory and experiment* **2008,** P10008 (2008).
3. Rabiner, L. R. A tutorial on hidden Markov models and selected applications in speech recognition. *Proceedings of the IEEE* **77,** 257–286 (1989).
4. Vidaurre, D., Smith, S. M. & Woolrich, M. W. Brain network dynamics are hierarchically organized in time. *Proceedings of the National Academy of Sciences* **114,** 12827–12832 (2017).
